# BNT162b vaccines are immunogenic and protect non-human primates against SARS-CoV-2

**DOI:** 10.1101/2020.12.11.421008

**Authors:** Annette B. Vogel, Isis Kanevsky, Ye Che, Kena A. Swanson, Alexander Muik, Mathias Vormehr, Lena M. Kranz, Kerstin C. Walzer, Stephanie Hein, Alptekin Güler, Jakob Loschko, Mohan S. Maddur, Ayuko Ota-Setlik, Kristin Tompkins, Journey Cole, Bonny G. Lui, Thomas Ziegenhals, Arianne Plaschke, David Eisel, Sarah C. Dany, Stephanie Fesser, Stephanie Erbar, Ferdia Bates, Diana Schneider, Bernadette Jesionek, Bianca Sänger, Ann-Kathrin Wallisch, Yvonne Feuchter, Hanna Junginger, Stefanie A. Krumm, André P. Heinen, Petra Adams-Quack, Julia Schlereth, Stefan Schille, Christoph Kröner, Ramón de la Caridad Güimil Garcia, Thomas Hiller, Leyla Fischer, Rani S. Sellers, Shambhunath Choudhary, Olga Gonzalez, Fulvia Vascotto, Matthew R. Gutman, Jane A. Fontenot, Shannan Hall-Ursone, Kathleen Brasky, Matthew C. Griffor, Seungil Han, Andreas A.H. Su, Joshua A. Lees, Nicole L. Nedoma, Ellene H. Mashalidis, Parag V. Sahasrabudhe, Charles Y. Tan, Danka Pavliakova, Guy Singh, Camila Fontes-Garfias, Michael Pride, Ingrid L. Scully, Tara Ciolino, Jennifer Obregon, Michal Gazi, Ricardo Carrion, Kendra J. Alfson, Warren V. Kalina, Deepak Kaushal, Pei-Yong Shi, Thorsten Klamp, Corinna Rosenbaum, Andreas N. Kuhn, Özlem Türeci, Philip R. Dormitzer, Kathrin U. Jansen, Ugur Sahin

## Abstract

A safe and effective vaccine against COVID-19 is urgently needed in quantities sufficient to immunise large populations. We report the preclinical development of two BNT162b vaccine candidates, which contain lipid-nanoparticle (LNP) formulated nucleoside-modified mRNA encoding SARS-CoV-2 spike glycoprotein-derived immunogens. BNT162b1 encodes a soluble, secreted, trimerised receptor-binding domain (RBD-foldon). BNT162b2 encodes the full-length transmembrane spike glycoprotein, locked in its prefusion conformation (P2 S). The flexibly tethered RBDs of the RBD-foldon bind ACE2 with high avidity. Approximately 20% of the P 2S trimers are in the two-RBD ‘down,’ one-RBD ‘up’ state. In mice, one intramuscular dose of either candidate elicits a dose-dependent antibody response with high virus-entry inhibition titres and strong TH1 CD4^+^ and IFNγ^+^ CD8^+^ T-cell responses. Prime/boost vaccination of rhesus macaques with BNT162b candidates elicits SARS-CoV-2 neutralising geometric mean titres 8.2 to 18.2 times that of a SARS-CoV-2 convalescent human serum panel. The vaccine candidates protect macaques from SARS-CoV-2 challenge, with BNT162b2 protecting the lower respiratory tract from the presence of viral RNA and with no evidence of disease enhancement. Both candidates are being evaluated in phase 1 trials in Germany and the United States. BNT162b2 is being evaluated in an ongoing global, pivotal Phase 2/3 trial (NCT04380701, NCT04368728).

## Introduction

Due to the shattering impact of the coronavirus disease 2019 (COVID-19) pandemic on human health and society, multiple collaborative research programs have been launched, generating new insights and progress in vaccine development. Soon after emerging in December 2019, the severe acute respiratory syndrome coronavirus-2 (SARS-CoV-2) was identified as a β-coronavirus with high sequence similarity to bat-derived SARS-like coronaviruses^1, 2^. Fast pandemic vaccine availability is critical, and the rapid globalised response is mirrored by the upload of over 212,000 viral genome sequences as of November 23, 2020, to GISAID (Global Initiative on Sharing All Influenza Data).

The trimeric spike glycoprotein (S) of SARS-CoV-2 is a key target for virus neutralising antibodies^3^ and the prime candidate for vaccine development. S binds its cellular receptor, angiotensin converting enzyme 2 (ACE2), through a receptor-binding domain (RBD), which is part of S1, its N-terminal furin cleavage fragment^4, 5^. On S, the RBDs have ‘up’ positions, in which the receptor binding sites and their dense cluster of neutralising epitopes are exposed, and ‘down’ positions, in which the receptor binding sites are buried, but some S neutralising epitopes on and off the RBDs remain available^6–9^. S rearranges to translocate the virus into cells by membrane fusion^6, 10^. The C-terminal furin cleavage fragment, S2, contains the fusion machinery^11^.

Messenger RNA technology allows versatile vaccine antigen design and highly scalable, fast manufacturing. With efficient lipid-nanoparticle (LNP) formulation processes, RNA vaccines are highly suited to rapid development and pandemic supply^12, 13^. RNA generated from DNA templates by a highly productive, cell-free *in vitro* transcription process is molecularly well defined and free of animal-origin materials. Here, we report the preclinical development of the LNP formulated N^1^-methyl-pseudouridine (m1Ψ) nucleoside-modified mRNA (modRNA) BNT162b vaccine candidates that encode SARS-CoV-2 S-derived immunogens (Fig. 1a). The m1Ψ-modification dampens innate immune sensing and, together with optimised non-coding sequence elements, increases efficiency of RNA translation *in vivo*^13–15^. Vaccines based on modRNA have proven immunogenic for several viral targets^16, 17^.

**Figure 1.**
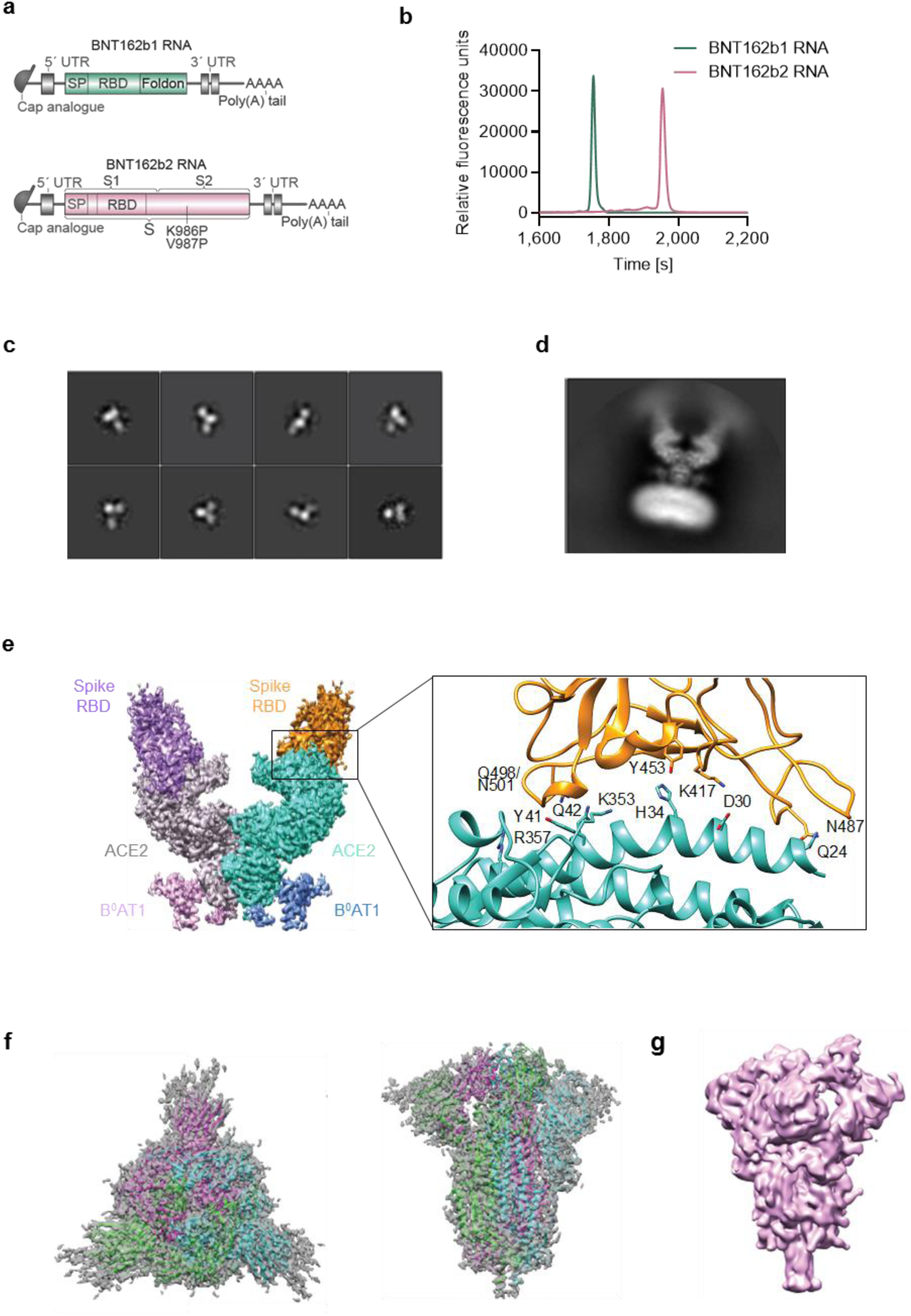
Vaccine design and characterisation of the expressed antigens. **a**, Structure of BNT162b RNAs. UTR, untranslated region; SP, signal peptide; RBD, receptor-binding domain; S1 and S2, N-terminal and C-terminal furin cleavage fragments, respectively; S, SARS-CoV-2 S glycoprotein. Proline mutations K986P and V897P are indicated. **b**, Liquid capillary electropherograms of both *in vitro* transcribed BNT162b RNAs. **c**, Representative 2D class averages from EM of negatively stained RBD-foldon trimers. Box edge: 37 nm. **d**, 2D class average from cryo-EM of the ACE2/B^0^AT1/RBD-foldon trimer complex. Long box edge: 39.2 nm. Peripheral to the relatively well-defined density of each RBD domain bound to ACE2, there is diffuse density attributed to the remainder of the flexibly tethered RBD-foldon trimer. A detergent micelle forms the density at the end of the complex opposite the RBD-foldon. **e**, Density map of the ACE2/B^0^AT1/RBD-foldon trimer complex at 3.24 Å after focused refinement of the ACE2 extracellular domain bound to a RBD monomer. Surface colour-coding by subunit. The ribbon model refined to the density shows the RBD-ACE2 binding interface, with residues potentially mediating polar interactions labeled. **f**, 3.29 Å cryo-EM map of P2 S, with fitted and refined atomic model, viewed down the three-fold axis toward the membrane (left) and viewed perpendicular to the three-fold axis (right). Coloured by protomer. **g,** Mass density map of TwinStrep-tagged P2 S produced by 3D classification of images extracted from cryo-EM micrographs with no symmetry averaging, showing the class in the one RBD ‘up’, two RBD ‘down’ position.

Both BNT162b vaccines are being evaluated in phase 1 clinical trials in the US (NCT04368728) and Germany (NCT04380701, EudraCT: 2020-001038-36); BNT162b2 is being evaluated in a pivotal, global, phase 2/3 safety and efficacy study^18–20^.

## Results

BNT162b1 RNA encodes the RBD with the SARS-CoV-2 S signal peptide (SP) fused to its N-terminus to enable ER translocation and secretion and with the trimerisation domain (foldon) of T4 fibritin^21^ fused to its C-terminus for multimeric display; BNT162b2 RNA encodes full-length S, stabilised in the prefusion conformation by the mutation of residues 986 and 987 to proline (P2 S; Fig. 1a)^7, 22, 23^. Both RNAs have single, sharp microfluidic capillary electrophoresis profiles, consistent with their calculated lengths, indicating high purity and integrity (Fig. 1b). Robust expression of RBD-foldon or P2 S was detectable by flow cytometry upon transfection of HEK293T cells with BNT162b1 RNA or BNT162b2 RNA, respectively, formulated as LNPs or mixed with a transfection reagent (Extended Data Fig. 1a). In transfected cells, BNT162b1-encoded RBD and BNT162b2-encoded P2 S localised to the secretory pathway as shown by immunofluorescence microscopy (Extended Data Fig. 1b). A main band of RBD-containing protein with an apparent MW >75 kDa was detected in the medium of BNT162b1 RNA-transfected cells (together with lesser quantities of a faster migrating species) by western blot under denaturing and non-denaturing conditions, consistent with secretion of trimeric RBD-foldon (predicted MW 88.4 kD; Extended Data Fig. 1c).

For further structural characterisation, the RBD-foldon and P2 S antigens were expressed from DNA corresponding to the RNA coding sequences. The RBD-foldon was purified from the medium of transfected Expi293F cells by affinity capture with the ACE2-peptidase domain (PD) immobilised on agarose beads, leaving little residual RBD-foldon uncaptured from the medium. Evidence that the RBD-foldon has three RBDs flexibly tethered to a central hub was obtained by electron microscopy (EM), which revealed a variety of conformations (Fig. 1c). The trimerised RBD bound to the human ACE2 peptidase domain (PD) with a KD of <5 pM, which is 1,000-fold the reported KD of 5 nM for monomeric RBD and consistent with the avidity effect of multivalent binding enabled by the flexible tethering (Extended Data Fig. 1d). Although the flexibility of the RBD-foldon precluded direct structural analysis at high resolution, one RBD per trimer could be immobilised by binding to a complex of ACE2 and the B^0^AT1 neutral amino acid transporter, which ACE2 chaperones, when that complex was in the previously reported closed conformation (Fig. 1d)^5^. The size and symmetry of the RBD-foldon/ACE2/B^0^AT1 ternary complex aided image reconstruction by electron cryomicroscopy (cryo-EM), and the structure of the RBD in the complex was determined to 3.24 Å resolution (Fig. 1e, Extended Data Table 1 and Supplementary Fig. 2). One copy of the RBD was resolved for each bound trimer. The binding interface between the resolved RBD and the ACE2 extracellular domain was fitted to a previously reported structure and showed good agreement^4^. The high avidity binding to ACE2 and well-resolved structure in complex with ACE2 demonstrate that the recombinant RBD-foldon authentically presents the ACE2 binding site targeted by many SARS-CoV-2 neutralising antibodies^8, 24^.

The trimeric P2 S was affinity purified from detergent solubilised protein via the C-terminal TwinStrep tag. P2 S bound the human ACE2-PD and a human anti-RBD neutralising antibody B38 with high affinity (KD 1 nM for each, Extended Data Fig. 1e, f)^25^. Structural analysis by cryo-EM produced a 3.29 Å nominal resolution mass density map, into which a previously published atomic model^7^ was fitted and rebuilt (Fig. 1f; Extended Data Fig. 2a, b and Table 1). The rebuilt model showed good agreement with reported structures of prefusion full-length wild type S and its ectodomain with P2 mutations^6, 7^. Three-dimensional classification of the dataset showed a class of particles that was in the one RBD ‘up’ (accessible for receptor binding), two RBD ‘down’ (closed) conformation and represented 20.4% of the trimeric molecules (Fig. 1g, Extended Data Fig. 2c). The remainder were in the all RBD ‘down’ conformation. The RBD in the ‘up’ conformation was less well resolved than other parts of the structure, suggesting conformational flexibility and a dynamic equilibrium between RBD ‘up’ and RBD ‘down’ states, as also suggested by others^6, 26^. The binding and structural analyses indicate that the BNT162b2 RNA sequence encodes a recombinant P2 S that can authentically present the ACE2 binding site and other epitopes targeted by SARS-CoV-2 neutralising antibodies.

To study vaccine immunogenicity, B- and T-cell responses were characterised in a series of experiments in BALB/c mice after a single intramuscular (IM) immunisation with 0.2, 1, or 5 µg of BNT162b vaccines, or buffer control. One immunisation with either candidate induced high dose level-dependent RBD- and S1-binding serum IgG titres (Fig, 2a, b; Extended Data Fig. 3a-c), which increased more steeply for BNT162b2. On day 28 after one immunisation with 5 µg BNT162b1 or BNT162b2, RBD-binding geometric mean endpoint titres were 752,680 or 434,560, respectively. IgG elicited by either candidate had strong binding affinity for a recombinant RBD target antigen (geometric mean KD 717 pM for BNT162b1 and 993 pM for BNT162b2), with a low off-rate and a high on-rate (Fig. 2c). Serum samples from buffer-immunised control animals had no detectable RBD- or S1-specific IgG (Fig. 2a, b and Extended Data Fig. 3a-c), and neither did serum samples from animals immunised up to two times with equivalent LNP-formulated modRNA that encoded a SARS-CoV-2 irrelevant antigen (not shown).

**Figure 2.**
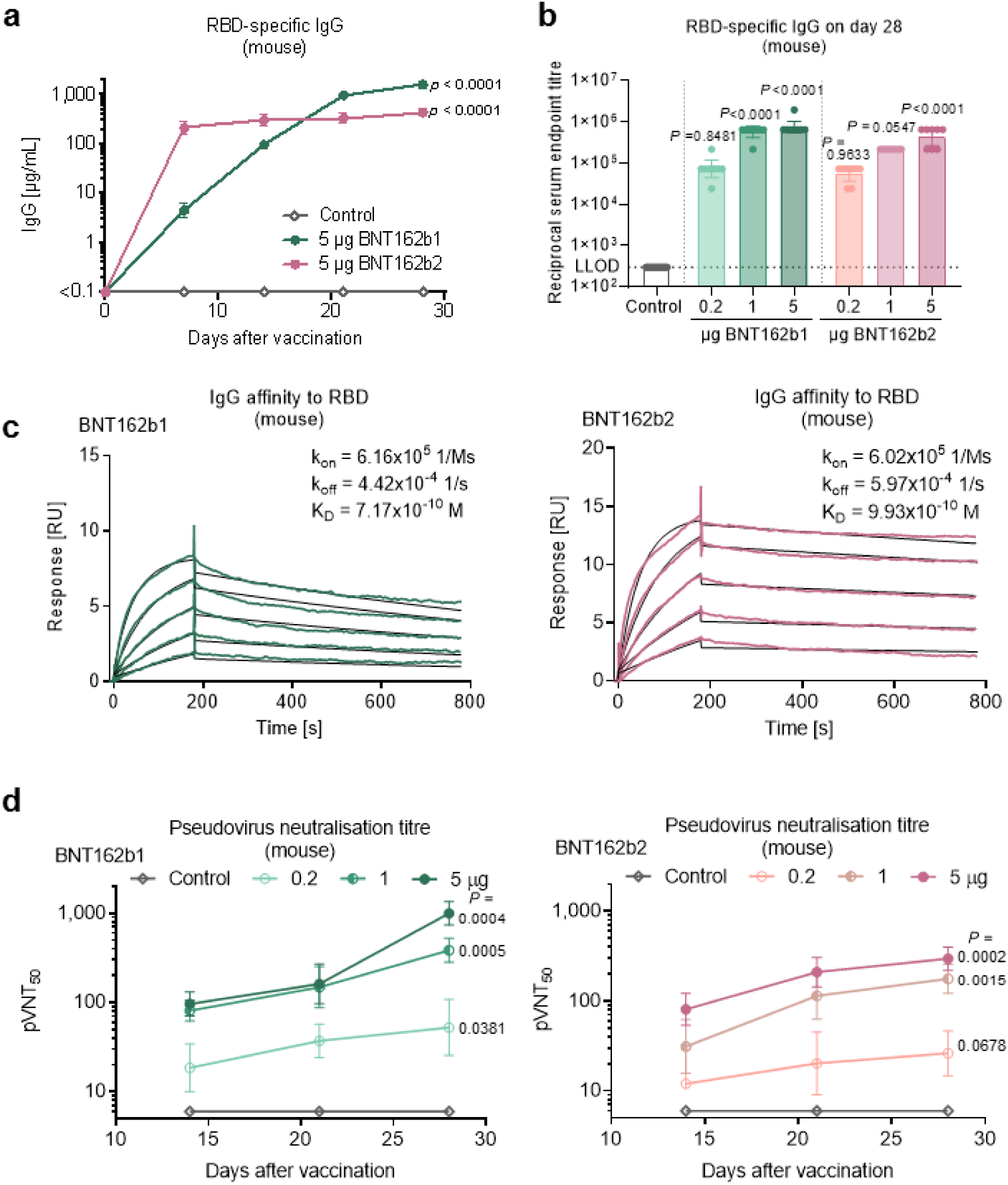
Mouse humoral immunogenicity. BALB/c mice (*n*=8) were immunised intramuscularly (IM) with a single dose of each BNT162b vaccine candidate or buffer control. Geometric mean of each group ± 95% confidence interval (CI) (a, b, d). Day 28 p-values compared to control (multiple comparison of mixed-effect analysis [a, d] and OneWay ANOVA [b], all using Dunnett’s multiple comparisons test) are provided. **a**, RBD-specific IgG levels in sera of mice immunised with 5 µg of BNT162b candidates, determined by ELISA. For day 0 values, a pre-screening of randomly selected animals was performed (*n*=4). For IgG levels with lower BNT162b doses and sera testing for detection of S1 see Extended Data Figure 3a, b. **b**, Reciprocal serum endpoint titres of RBD-specific IgG 28 days after immunisation. The horizontal dotted line indicates the lower limit of detection (LLOD). **c**, Representative surface plasmon resonance sensorgrams of the binding kinetics of His-tagged RBD to immobilised mouse IgG from serum drawn 28 days after immunisation with 5 µg of each BNT162b. Actual binding (in colour) and the best fit of the data to a 1:1 binding model (black) are depicted. For binding kinetics of same sera to His-tagged S1 see Extended Data Figure 3d. **d**, Pseudovirus-based VSV-SARS-CoV-2 50% neutralisation titres (pVNT50) in sera of mice immunised with BNT162b vaccine candidates. For number of infected cells per well with serum samples drawn 28 days after immunisation and titre correlation to a SARS-CoV-2 virus neutralisation assay see Extended Data Figure 3e-g.

Virus entry inhibition by BNT162b immunised mouse serum was measured with a vesicular stomatitis virus (VSV)-based SARS-CoV-2 pseudovirus neutralisation assay. Like the antigen-specific IgG geometric mean titres (GMTs), fifty percent pseudovirus neutralisation (pVNT50) GMTs increased steadily after immunisation with 5 μg of either candidate, reaching 1,056 for BNT162b1 and 296 for BNT162b2 on Day 28 after immunisation (Fig. 2d, Extended Data Fig. 3e, f). A random selection of samples was tested in a SARS-CoV-2 virus neutralisation assay, demonstrating strong correlation of pseudovirus and SARS-CoV-2 neutralisation (Pearson correlation of 0.9479 between the tests (Extended Data Fig. 3g). In summary, each candidate induced a high functional antibody response in mice, with BNT162b1 inducing higher titres after one immunisation.

Characterisation of antigen-specific splenic T-cell responses in mice 12 and 28 days after BNT162b vaccine immunisation revealed a high fraction of CD4^+^ and CD8^+^ T cells that produced IFNγ and CD8^+^ cells that produced IL-2, as shown by enzyme linked immunospot assay (ELISpot) or intracellular cytokine staining (ICS) flow cytometry analysis after *ex vivo* restimulation with a full-length S peptide pool (Fig. 3a-c). Total splenocytes harvested on Day 28 and re-stimulated with the full-length S peptide pool secreted high levels of the TH1 cytokines IL-2 or IFNγ and minute or undetectable levels of the TH2 cytokines IL-4, IL-5 or IL-13, as measured in multiplex immunoassays (Fig. 3d). Overall, the patterns of CD4^+^ and CD8^+^ T-cell responses were similar for the two vaccine candidates, with a somewhat stronger IFNγ-producing CD8^+^ T-cell response in BNT162b2-immunised mice.

**Figure 3.**
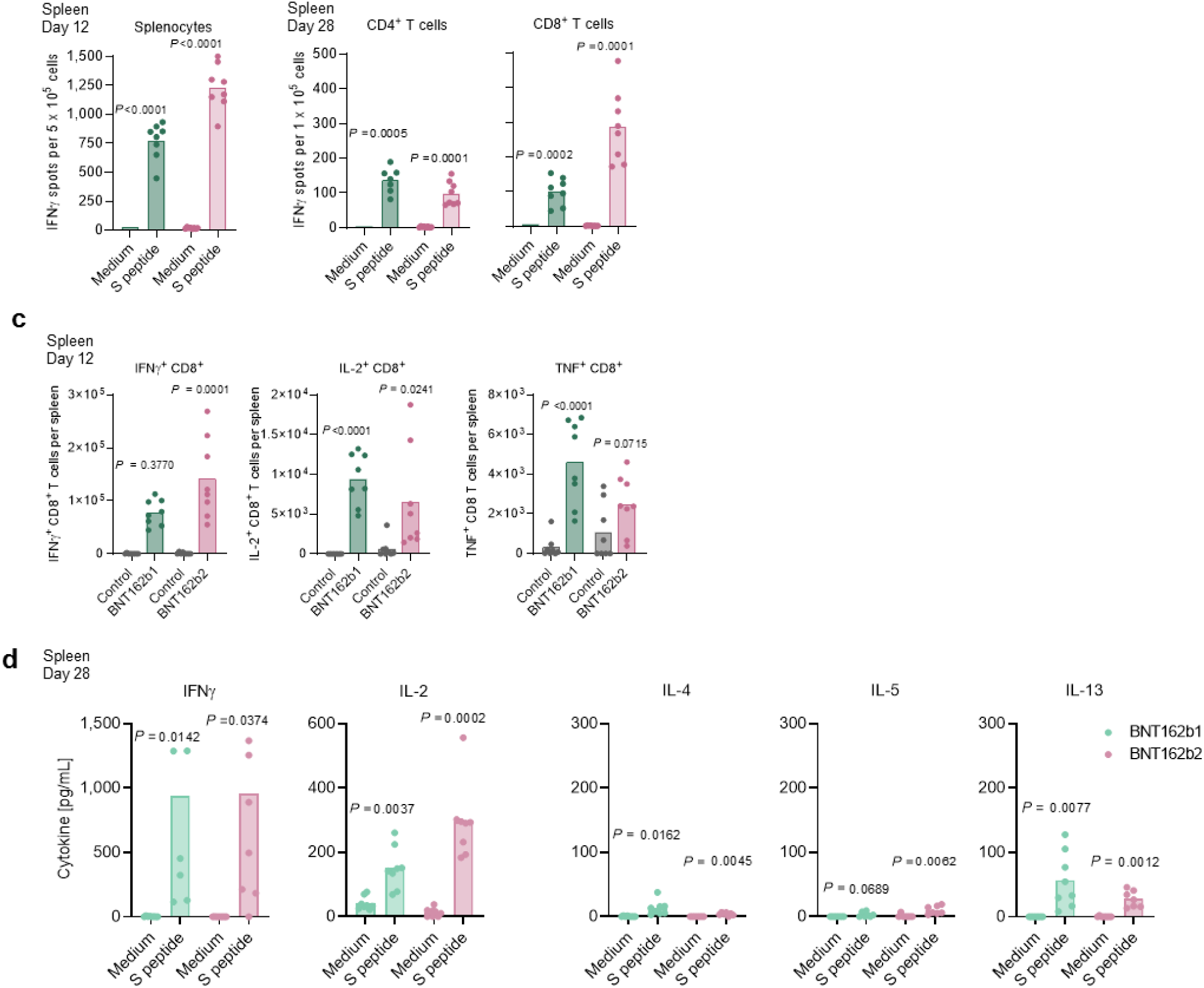
Mouse cellular immunogenicity. Splenocytes of BALB/c mice (*n*=8, unless stated otherwise) immunised IM with BNT162b vaccines were *ex vivo* re-stimulated with full-length S peptide mix (a-d) or cell culture medium (a, b, d). Symbols represent individual animals. Means of each group are shown, P-values compare immunised groups with the control (two-tailed paired t-test). **a**, IFNγ ELISpot of splenocytes after immunisation with 5 μg BNT162b vaccines. **b**, IFNγ ELISpot of splenic CD4^+^ or CD8^+^ T cells after immunisation with 1 µg BNT162b vaccines (BNT162b1: *n*=7 for CD4^+^ T cells, one outlier removed by Grubbs test, α=0.05). **c**, CD8^+^ T-cell specific cytokine release by splenocytes after immunisation with 5 µg BNT162b vaccines or buffer (Control), determined by flow cytometry. S-peptide specific responses are corrected for background (medium). **d**, Cytokine production by splenocytes after immunisation with 0.2 µg BNT162b1 or 1 µg BNT162b2, determined by bead-based multiplex analysis (BNT162b2: *n*=7 for IL-4, IL-5 and IL-13, one outlier removed by the ROUT method [Q=1%] for the S peptide stimulated samples).

Vaccine-induced effects on the proliferation and dynamics of immune cell populations were assessed in injection site draining lymph nodes (dLNs), to evaluate the principal immune-educated compartments for proficient T- and B-cell priming, as well as in blood and spleen, to evaluate systemic vaccine effects. Higher numbers of plasma cells, class switched IgG1- and IgG2a-positive B cells, and germinal center B cells were observed in dLNs, and higher numbers of class switched IgG1-positive and germinal centre B cells were observed in spleens of mice 12 days after immunisation with 5 μg of either vaccine as compared to control (Extended Data Fig. 4a, b). Vaccine-immunised mice had significantly fewer circulating B cells than control mice as measured in blood at Day 7 post-immunisation (Extended Data Fig. 4c), which may imply that B-cell homing to lymphoid compartments contributed to augmented B-cell counts in dLN and spleen.

The dLNs from BNT162b1- or BNT162b2-immunised mice also displayed significantly elevated counts of CD8^+^ and CD4^+^ T cells, which were most pronounced for T follicular helper (TFH) cells, including ICOS^+^ subsets that are essential for germinal centre formation (Extended Data Fig. 4a). Both BNT162b vaccines increased TFH cell counts in the spleen and blood, while an increase in circulating CD8^+^ T cells was only detected in BNT162b2-immunised mice (Extended Data Fig. 4b, c). In aggregate, these data indicate a strong induction of SARS-CoV-2 pseudovirus neutralisation titres and systemic CD8^+^ and TH1-driven CD4^+^ T-cell responses by both modRNA vaccine candidates, with a somewhat more pronounced cellular response to BNT162b2.

To assess the immunogenicity of BNT162b1 and BNT162b2 in non-human primates, groups of six male, 2-4 year old rhesus macaques were immunised IM with 30 or 100 μg of BNT162b1, BNT162b2, or saline control on Days 0 and 21. RBD-binding IgG was readily detectable by Day 14 after Dose 1, and levels increased further 7 days after Dose 2 (Day 28; Fig. 4a). On Day 28, geometric mean RBD-binding IgG concentrations (GMCs) were 20,962 units (U)/mL (30 μg dose level) and 48,575 U/mL (100 μg dose level) for BNT162b1 and 23,781 U/mL (30 μg dose level) and 26,170 U/mL (100 μg dose level) for BNT162b2. For comparison, the RBD-binding IgG GMC of a panel of 38 SARS-CoV-2 convalescent human sera (HCS) was 602 U/mL, lower than the GMC of immunised rhesus macaques after one or two doses.

**Figure 4.**
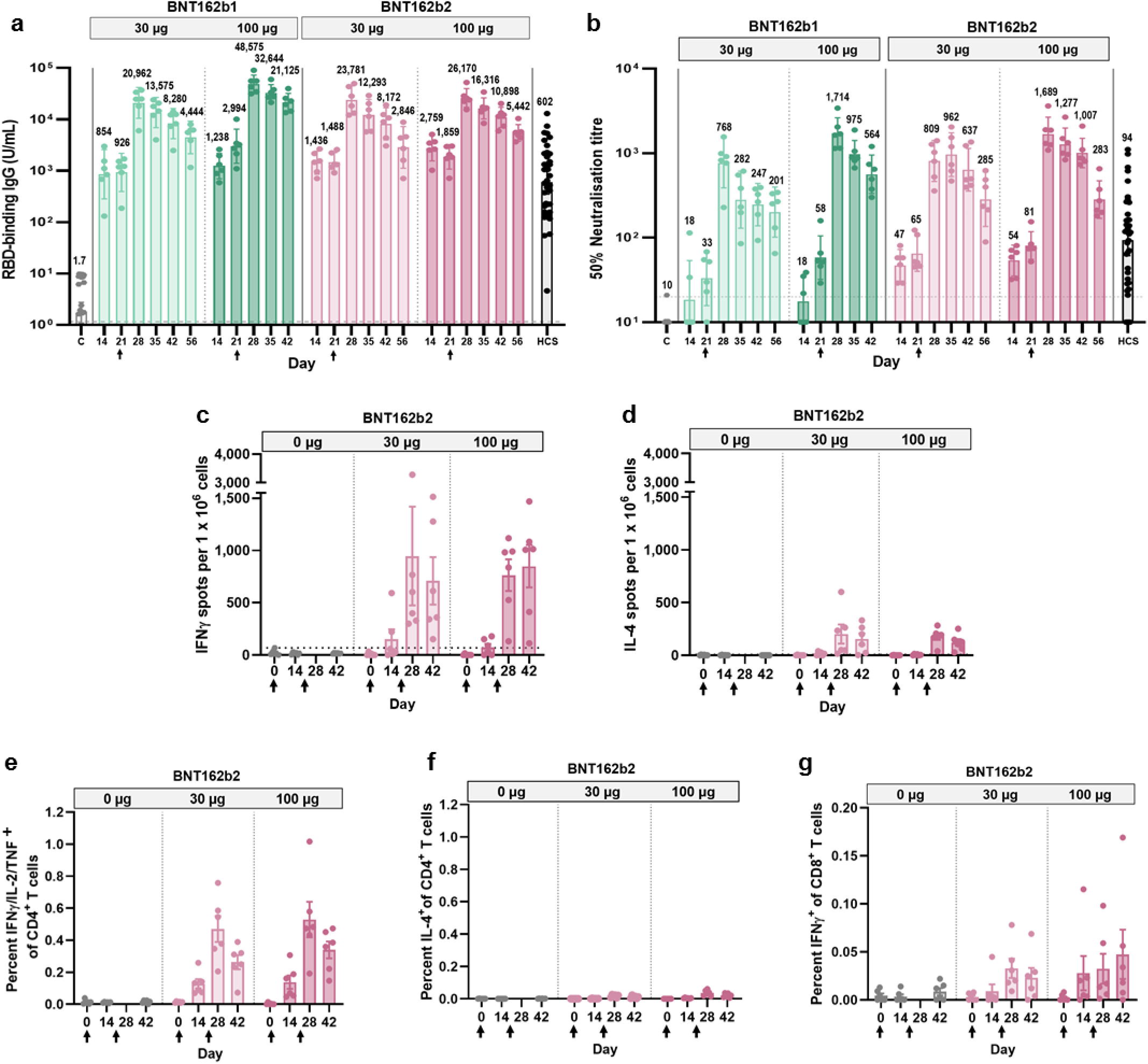
Rhesus macaque immunogenicity. Male rhesus macaques, 2-4 years of age, were immunised on Days 0 and 21 (arrows below the x-axis indicate the days of the second immunisation) with 30 µg or 100 µg BNT162b vaccines (*n*=6 each). Additional rhesus macaques received saline (C; *n*=9). Human convalescent sera (HCS) were obtained from SARS-CoV-2-infected patients at least 14 days after PCR-confirmed diagnosis and at a time when acute COVID-19 symptoms had resolved (*n*=38). The HCS panel is a benchmark for serology studies in this and other manuscripts. **a**, Concentrations, in arbitrary units, of IgG binding recombinant SARS-CoV-2 RBD (LLOD = 1.72 U/mL). **b**, SARS-CoV-2 50% virus neutralisation titres (VNT50, LLOD = 20). **c-g**, PBMCs collected on Days 0, 14, 28 and 42 were *ex vivo* re-stimulated with full-length S peptide mix. **c,** IFNγ ELISpot. **d,** IL-4 ELISpot. **e**, S-specific CD4^+^ T-cell IFNγ, IL-2, or TNFα release by flow cytometry (LLOD = 0.04). **f**, S-specific CD4^+^ T-cell IL-4 release by flow cytometry (LLOD = 0.05). **g**, CD8^+^ T-cell IFNγ release by flow cytometry (LLOD = 0.03). Heights of bars indicate the geometric (a-b) or arithmetic (c-g) means for each group, with values written above bars (a-b). Whiskers indicate 95% confidence intervals (CI’s; a-b) or standard errors of means (SEMs; c-g). Each symbol represents one animal. Horizontal dashed lines mark LLODs. For serology and ELISpot data (a-d) but not for flow cytometry data (e-g), values below the LLOD were set to ½ the LLOD. Arrows below the x-axis indicate the days of Doses 1 and 2.

Fifty percent virus neutralisation GMTs, measured by a SARS-CoV-2 neutralisation assay^27^ (not a pseudovirus neutralisation assay), were detectable in the sera of most BNT162b1-immunised rhesus macaques by Day 21 after Dose 1 and in all BNT162b2-immunised macaques by Day 14 after Dose 1 (Fig. 4b). There was a strong boosting effect, with comparable GMTs elicited by BNT162b1 (768 for 30 μg and 1,714 for 100 μg) or BNT162b2 (962 for 30 μg or 1,689 for 100 μg), measured in sera drawn 7 or 14 days after Dose 2. For BNT162b2, sera were available up to Day 56 after Dose 1 (28 days after Dose 2), and robust GMTs of 285 for 30 μg and 283 for 100 μg dose levels persisted to that time point. For comparison, the neutralisation GMT of the human convalescent serum was 94, substantially lower than the GMTs of rhesus macaque sera drawn 21 or 35 days after Dose 2.

S-specific T-cell responses of the BNT162b2- or saline-immunised rhesus macaques were analysed using peripheral blood mononuclear cells (PBMCs) collected before immunisation and at the times indicated after Doses 1 and 2. ELISpot demonstrated strong IFNγ but minimal IL-4 responses after Dose 2 (Fig. 4c, d, and Extended Data Fig. 5). ICS confirmed that BNT162b2 elicited a high frequency of CD4^+^ T cells that produced IFNγ, IL-2, or TNF but a low frequency of CD4^+^ T cells that produced IL-4, indicating a TH1-biased response (Fig. 4e, f). ICS also demonstrated that BNT162b2 elicited circulating S-specific CD8^+^ T cells that produced IFNγ (Fig. 4g).

Forty-one to fifty-five days after Dose 2, 6 of the 2-4 year old rhesus macaques that had been immunised with 100 μg BNT162b1 and 6 that had been immunised with 100 μg BNT162b2 were challenged with 1.05 × 10^6^ plaque forming units of SARS-CoV-2 (strain USA-WA1/2020), split equally between intranasal and intratracheal routes, as previously described (Extended Data Fig. 6, Extended Data Table 2)^28^. In addition, nine age-matched macaques (controls) that had been mock-immunised with saline received the same SARS-CoV-2 challenge, and 6 age-matched macaques (sentinels), 3 of which had been immunised with 30 µg BNT162b2, were mock-challenged with cell culture medium. Nasal, oropharyngeal (OP), and rectal swabs were collected, and bronchoalveolar lavage (BAL) was performed at the times indicated (Extended Data Table 2). Samples were tested for SARS-CoV-2 RNA (genomic RNA and subgenomic transcripts) by reverse-transcription quantitative polymerase chain reaction (RT-qPCR; Fig. 5a,b). All personnel performing clinical, radiological, histopathological, or RT-qPCR evaluations were blinded to the group assignments of the macaques.

**Figure 5.**
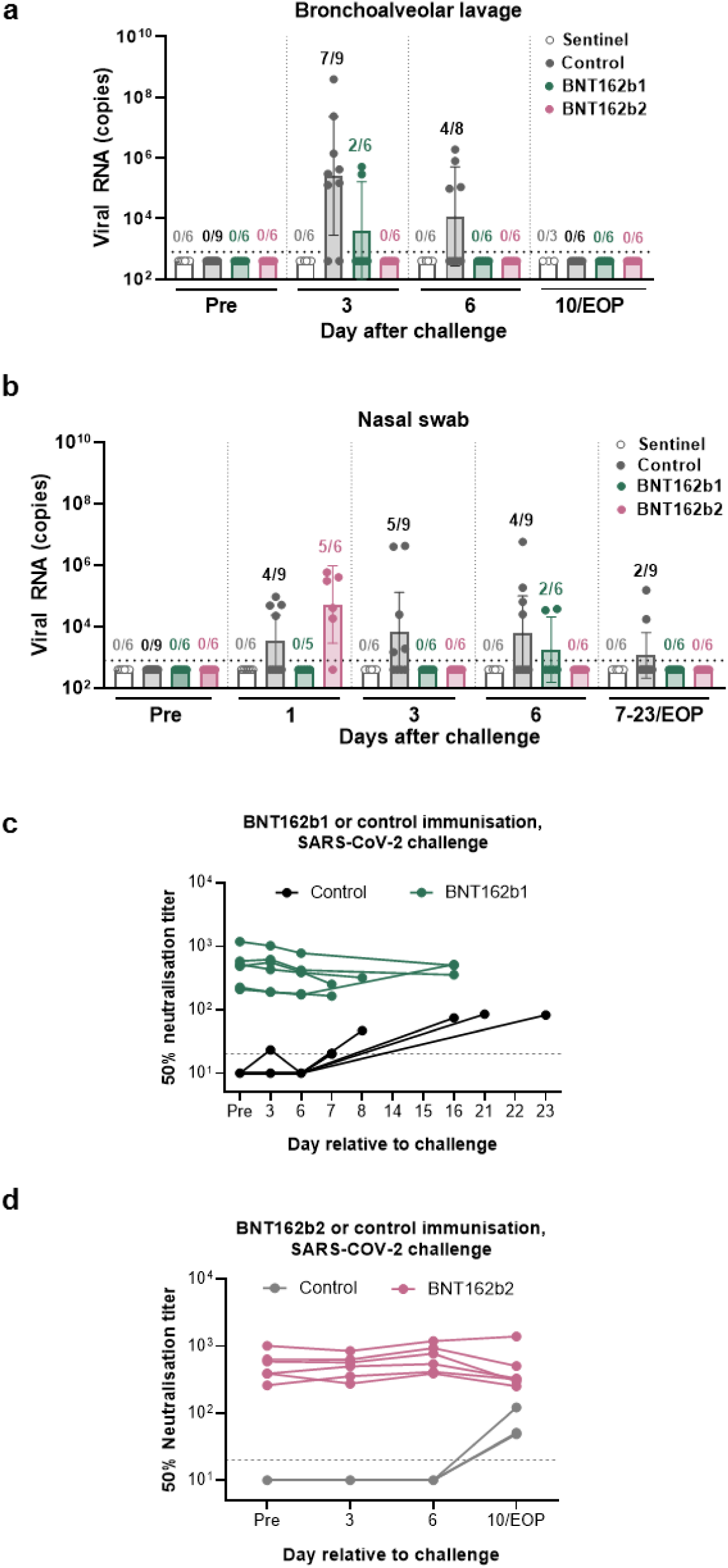
Virological and serological evidence of protection of rhesus macaques from challenge with infectious SARS-CoV-2. Rhesus macaques immunised with 100 µg of BNT162b1 or BNT162b2 (*n*=6 each) or mock immunised with saline challenge (Control, *n*=9) were challenged with 1.05 × 10^6^ total plaque forming units (PFU) of SARS-CoV-2 split equally between the intranasal (IN) and intratracheal (IT) routes. Additional macaques (Sentinel, *n*=6) were mock-challenged with cell culture medium. Macaque assignments to cohorts and schedules of immunisation, challenge, and sample collection are provided in Extended Data Fig. 6 and Extended Data Table 2. Viral RNA levels were detected by RT-qPCR. **a,** Viral RNA in bronchoalveolar lavage (BAL) fluid. **b**, Viral RNA in nasal swabs. Symbols represent individual animals. Ratios above bars indicate the number of viral RNA positive animals among all animals in a group with evaluable samples. Heights of bars indicate geometric mean viral RNA copies; whiskers indicate geometric standard deviations. Each symbol represents one animal. Dotted lines indicate the lower limit of detection (LLOD). Values below the LLOD were set to ½ the LLOD. The statistical significance by a non-parametric test (Friedman’s test) of differences in viral RNA detection after challenge between 6 BNT162b1-immunised and 6 mock-immunised animals (challenge cohorts 1 and 2) was p = 0.015 for BAL fluid and p = 0.005 for nasal swab; between 6 BNT162b2-immunised animals and 3 mock-immunised animals (challenge cohort 3), the statistical significance was p = 0.001 for BAL fluid and p = 0.262 for nasal swabs. Serum samples were assayed for SARS-CoV-2 50% neutralisation titres (VNT50). **c**, BNT162b1-immunised macaques and Controls (challenge cohorts 1 and 2). **d**, BNT162b2-immunised macaques and Controls (challenge cohort 3). Symbols represent individual animal titres. Horizontal dashed lines indicate the LLOQ of 20.

Viral RNA was detected in BAL fluid from 7 of the 9 control macaques on Day 3, from 4 of 8 on Day 6 after challenge (with 1 indeterminant result), and from none of the 6 that underwent BAL at the end of project (EOP, Days 7-23 after challenge; Fig. 5a). Viral RNA was detected in the BAL fluid of 2 of 6 BNT162b1-immunised macaques on day 3 after challenge and from none thereafter. At no time point sampled was viral RNA detected in BAL fluid from the BNT162b2-immunised and SARS-CoV-2 challenged macaques.

In nasal swabs obtained on the day after challenge, viral RNA was detected from control-immunised macaques (4 of 9) and BNT162b2-immunised macaques (5 of 6) but not from BNT162b1-immunised macaques (Fig. 5b). In subsequent nasal swabs, viral RNA was detected from some of the control-immunised macaques on each sampling (5 of 9 on Day 3, 4 of 9 on Day 6, and 2 of 9 on Days 7-23), from some BNT162b1-immunised macaques on only 1 sampling (2 of 6 on Day 6), and from none of the BNT162b2-immunised macaques on any sampling. Similar patterns were seen in OP and rectal swabs, with viral RNA more often detected in control-immunised macaques than in BNT162b1- or BNT162b2-immunised macaques and with more persistence of viral RNA in rectal swabs than in OP swabs (Extended Data Fig. 7a, b).

At the time of challenge, SARS-CoV-2 neutralising titres ranged from 208 to 1,185 in the BNT162b1-immunised animals and from 260 to 1,004 in the BNT162b2-immunised animals. Neutralising titres were below the limit of detection in the control animals (Fig. 5c, d). The control animals responded to infectious virus challenge with an increase in SARS-CoV-2 neutralising titres, consistent with an immune response to viral infection. However, there was no trend toward increasing SARS-CoV-2 neutralising titres in response to viral challenge in the BNT162b1-immunised or BNT162b2-immunised animals, consistent with immunisation suppressing SARS-CoV-2 infection. The maximum SARS-CoV-2 neutralising titre elicited by virus challenge of control rhesus macaques remained below 150 through the time of necropsy, whereas all immunised animals maintained neutralising titres greater than 150 throughout the challenge experiment.

None of the challenged animals, whether immunised or not, showed clinical signs of illness (Extended Data Fig. 8a-d). Radiographic abnormalities were generally minimal or mild and were not consistently associated with viral challenge (Extended Data Fig. 9a, b). Histopathology of necropsy specimens obtained 7-8 days after challenge revealed localised areas of pulmonary inflammation that were limited in extent even in the control animals challenged after mock immunisation with saline (Extended Data Fig. 10). We conclude that the 2-4 year old male rhesus macaque challenge model is primarily a SARS-CoV-2 infection model rather than a COVID-19 disease model.

## Discussion

We demonstrate that BNT162b1 or BNT162b2, LNP-formulated, m1Ψ nucleoside-modified mRNAs that encode secreted, trimerised SARS-CoV-2 RBD or prefusion-stabilised S, respectively, induce strong antigen-specific immune responses in mice and rhesus macaques. The RBD-foldon coding sequence directs the expression and secretion of a flexible, trimeric protein that binds ACE2 with high affinity and has structurally intact ACE2 receptor binding sites. Protein expressed from DNA with the BNT162b2-encoded P2 S amino acid sequence was confirmed to be in the prefusion conformation by cryo-EM. This analysis showed that the antigenically important RBD can assume the ‘up’ conformation, with the receptor binding site, rich in neutralising epitopes, accessible in a proportion of the molecules^24^. The alternative states observed likely reflect a dynamic equilibrium between RBD ‘up’ and ‘down’ positions^7, 26^. Binding of expressed and purified P2 S to ACE2 and a neutralising monoclonal antibody further demonstrates its conformational and antigenic integrity.

In mice, a single sub-microgram immunisation with either BNT162b candidate rapidly induced high antibody titres that inhibited pseudovirus entry in the range of or above recently reported neutralising titres elicited by other SARS-CoV-2 vaccine candidates^29, 30^. The candidates also induced strong TFH and TH1 type CD4^+^ T-cell responses, the latter thought to be a more general effect of LNP-formulated modRNA vaccines against SARS-CoV-2^31^. Both CD4^+^ T-cell types are known to support antigen-specific antibody generation and maturation. In some animal models of respiratory virus infection, a TH2 type CD4^+^ T-cell response has been associated with vaccine-associated enhanced respiratory disease^32, 33^. Therefore, a TH1 type response to immunisation is preferred, as it may reduce the theoretical risk of enhanced pulmonary disease during subsequent viral infection. Immunisation with the vaccine candidates triggered redistribution of B cells from the blood to lymphoid tissues, where antigen presentation occurs. In humans, TFH cells in the circulation after vaccination with a VSV-vectored Ebola vaccine candidate have been correlated with a high frequency of antigen-specific antibodies^34^. After vaccination of mice with BNT162b1 or BNT162b2, high numbers of TFH were present in both blood and LN, a potential correlate for the generation of a strong adaptive B-cell response in germinal centres. In addition to eliciting favourable CD4^+^ T-cell responses, both BNT162b1 and BNT162b2 elicit CD8^+^ T-cell responses in mice, with BNT162b2 appearing to be somewhat more efficient at eliciting antigen-specific cytotoxic IFNγ CD8^+^ T cells.

BNT162b1 and BNT162b2 elicit immune profiles in rhesus macaques similar to those observed in mice. Seven days after Dose 2 of 100 µg administered to macaques, during the expansion phase of the antibody response, neutralising GMTs elicited by either candidate reached approximately 18-times the GMT of a human SARS-CoV-2 convalescent serum panel. Neutralising GMTs declined by Day 56 (35 days after Dose 2), consistent with the contraction phase, but remained well above the GMT of the panel. The duration of the study was not long enough to assess the rate of decline during the plateau phase of the antibody response. As it had in mice, BNT162b2 elicted a strongly TH1-biased CD4^+^ T-cell response and IFNγ^+^ CD8^+^ T-cell response in rhesus macaques.

Limitation and clearance of virus infections is promoted by the interplay of neutralising antibodies that eliminate infectious particles with CD8^+^ T cells that target intracellular virus reservoirs. CD8^+^ T cells may also reduce the influx of monocytes into infected lung tissue, which can be associated with undesirable IL-6 and TNF production and impaired antigen presentation^35, 36^. The responses elicited by the vaccine candidates reflect a pattern favourable for vaccine safety and efficacy, providing added reassurance for clinical translation^37^. The contributions of the individual immune effector systems to human protection from SARS-CoV-2 are not yet understood. Therefore, it appears prudent to develop COVID-19 vaccines that enlist concomitant cognate B cells, CD4^+^ T cells, and CD8^+^ T-cell responses.

Both candidates protected 2-4 year old rhesus macaques from infectious SARS-CoV-2 challenge, with reduced detection of viral RNA in immunised animals compared to those that received saline. Immunisation with BNT162b2 provided particularly strong RT-qPCR evidence for lower respiratory tract protection, as demonstrated by the absence of detectable SARS-CoV-2 RNA in serial BAL samples obtained starting 3 days after challenge. The lack of serological response to the SARS-CoV-2 challenge in BNT162b1- or BNT162b2-immunised macaques, despite a neutralising response to challenge in control-immunised macaques, suggests suppression of infection by the vaccine candidates. Clinical signs of disease were absent, and radiological and pathological abnormalities were generally mild after challenge. There was no evidence of vaccine-mediated enhancement of viral replication, disease, or pathology.

The selection of BNT162b2 over BNT162b1 for further clinical testing was largely driven by greater tolerability of BNT162b2 with comparable immunogenicity in clinical trials^20^ and the broader range and MHC-diversity of T-cell epitopes on the much larger full-length spike. A global, pivotal, phase 3 safety and efficacy study of immunisation with BNT162b2 (NCT04368728) is ongoing and may answer those open questions that cannot be addressed by preclinical models.

## Materials and Methods

### Ethics statement

All mouse studies were performed at BioNTech SE, and protocols were approved by the local authorities (local welfare committee), conducted according to Federation of European Laboratory Animal Science Associations recommendations and in compliance with the German Animal Welfare Act and Directive 2010/63/EU. Only animals with an unobjectionable health status were selected for testing procedures.

Immunisations for the non-human primate (NHP) study were performed at the University of Louisiana at Lafayette-New Iberia Research Centre (NIRC), which is accredited by the Association for Assessment and Accreditation of Laboratory Animal Care (AAALAC, Animal Assurance #: 000452). The work was in accordance with USDA Animal Welfare Act and Regulations and the NIH Guidelines for Research Involving Recombinant DNA Molecules, and Biosafety in Microbiological and Biomedical Laboratories. All procedures performed on these animals were in accordance with regulations and established guidelines and were reviewed and approved by an Institutional Animal Care and Use Committee or through an ethical review process. Infectious SARS-CoV-2 challenge of NHPs following immunisation was performed at the Southwest National Primate Research Centre (SNPRC), Texas Biomedical Research Institute, which is also accredited by the Association for Assessment and Accreditation of Laboratory Animal Care (AAALAC, Animal Assurance #: 000246). Animal husbandry followed standards recommended by AAALAC International and the NIH Guide for the Care of Use of Laboratory Animals. This study was approved by the Texas Biomedical Research Institute Animal Care and Use Committee.

### Protein and peptide reagents

Purified recombinant SARS-CoV-2 RBD (Sino Biological) or trimeric S protein (Acro Biosystems) was used as a target for western blot, and the RBD tagged with a human Fc (Sino Biological) was used in ELISA to detect SARS-CoV-2 S-specific IgG. A recombinant SARS-CoV-2 RBD containing a C-terminal Avitag™ (Acro Biosystems) was used as a target antigen in Luminex immunoassays. Purified recombinant SARS-CoV-2 S1 including a histidine tag (Sino Biological) was used in ELISA to detect SARS-CoV-2 S-specific IgG in mice. Purified recombinant SARS-CoV-2 S1 and RBD with histidine tags (both Sino Biological) were used for surface plasmon resonance (SPR) spectroscopy. A peptide pool of 15-mer peptides overlapping by 11 amino acids covering the full length S protein was used for re-stimulation in ELISpot, cytokine profiling and intracellular cytokine staining followed by flow cytometry. An irrelevant peptide (SPSYVYHQF, derived from gp70 AH-1^38^) or a CMV peptide pool was used as control for ELISpot assays. All peptides were obtained from JPT Peptide Technologies.

### Human convalescent sera

Human COVID-19 convalescent sera (*n*=38) were drawn from donors 18-83 years of age at least 14 days after PCR-confirmed diagnosis and at a time when the participants were asymptomatic. Most serum donors had outpatient (35/38) or inpatient (1/38) COVID-19; two of thirty-eight had asymptomatic SARS-CoV-2 infections. Sera were obtained from Sanguine Biosciences (Sherman Oaks, CA), the MT group (Van Nuys, CA) and Pfizer Occupational Health and Wellness (Pearl River, NY) and were used across different studies as a reference benchmark panel^18–20^.

### Cell culture

Human embryonic kidney (HEK)293T and Vero 76 cells (both ATCC) were cultured in Dulbecco’s modified Eagle’s medium (DMEM) with GlutaMAX™ (Gibco) supplemented with 10% fetal bovine serum (FBS [Sigma-Aldrich]). Cell lines were tested for mycoplasma contamination after receipt, before expansion and cryopreservation. For studies including NHP samples, Vero 76 and Vero CCL81 cells (both ATCC) were cultured in DMEM (Gibco) containing 2% HyClone fetal bovine and 100 U/mL penicillium/streptomycin (Gibco). Expi293F™ cells were grown in Expi293™ media and transiently transfected using ExpiFectamine™293 (all from Thermo Fisher Scientific).

### *In vitro* transcription and purification of RNA

Antigens encoded by BNT162b vaccine candidates were designed on a background of S sequences from SARS-CoV-2 isolate Wuhan-Hu-1 (GenBank: MN908947.3). The DNA template for the BNT162b1 RNA is a DNA fragment encoding a fusion protein of the SARS-CoV-2 S signal peptide (SP, amino acids 1-16), the SARS-CoV-2 S RBD, and the T4 bacteriophage fibritin trimerisation motif^21^ (‘foldon’). The template for the BNT162b2 RNA is a DNA fragment encoding SARS-CoV-2 S (GenBank: MN908947) with K986P and V987P mutations. BNT162b1 and BNT162b2 DNA templates were cloned into a plasmid vector with backbone sequence elements (T7 promoter, 5′ and 3′ UTR, 100 nucleotide poly(A) tail) interrupted by a linker (A30LA70, 10 nucleotides) for improved RNA stability and translational efficiency^14, 39^. The DNA was purified, spectrophotometrically quantified, and *in vitro* transcribed by T7 RNA polymerase in the presence of a trinucleotide cap1 analogue ((m2^7,3’-O^)Gppp(m^2’-O^)ApG; TriLink) and with N^1^-methylpseudouridine-5’-triphosphate (m1ΨTP; Thermo Fisher Scientific) replacing uridine-5’-triphosphate (UTP)^40^. RNA was purified using magnetic particles^41^. RNA integrity was assessed by microfluidic capillary electrophoresis (Agilent Fragment Analyser), and the concentration, pH, osmolality, endotoxin level and bioburden of the solution were determined.

### Lipid-nanoparticle formulation of the RNA

Purified RNA was formulated into LNPs using an ethanolic lipid mixture of ionisable cationic lipid and transferred into an aqueous buffer system via diafiltration to yield an LNP composition similar to one previously described^42^. The vaccines candidates were stored at −70 to −80 °C at a concentration of 0.5 mg/mL.

### Transfection of HEK cells

HEK293T cells were transfected with 1 µg RiboJuice transfection reagent-mixed BNT162b1 RNA or BNT162b2 RNA or with the vaccine candidates BNT162b1 (LNP-formulated BNT162b1 RNA) or BNT162b2 (LNP-formulated BNT162b2 RNA) by incubation for 18 hours. Non-LNP formulated mRNA was diluted in Opti-MEM medium (Thermo Fisher Scientific) and mixed with the transfection reagent according to the manufacturer’s instructions (RiboJuice, Merck Millipore).

### Western blot analysis of size fractions of the medium of BNT162b1 RNA transfected cells

Medium from cultured HEK293T cells were collected. After 13-fold concentration via Vivaspin 20 centrifugal concentrators with a molecular weight cut off of 10 kDa, supernatants were applied to a preparative HiLoad^®^ 16/600 Superdex^®^ 200 pg column (both Sigma Aldrich). The column was run at 29.8 cm/h in phosphate buffered saline (PBS), and 500 µL fractions were collected (Supplementary Fig. 1). The gel filtration column was calibrated with well defined protein standards separated under identical conditions in a second run. Size fractioned FBS-free medium from BNT162b1 RNA-transfected HEK293T cells was analysed by denaturing (95° C) and non-denaturating (no-heating) PAGE using 4–15% Criterion™ TGX Stain-Free™ Gel (Bio-Rad) and western blot. Transfer to a nitrocellulose membrane (Bio-Rad) was performed using a semi-dry transfer system (Trans-Blot Turbo Transfer System, Bio-Rad). Blotted proteins were detected with a monoclonal antibody that recognizes SARS-CoV-2 S1 (SinoBiological) and a secondary anti-rabbit horse radish peroxidase (HRP)-conjugated antibody (Sigma Aldrich). Blots were developed with Clarity Western ECL Substrate (Bio-Rad) and imaged with a Fusion FX Imager (Vilber) using the Image Lab software version 6.0.

### Vaccine antigen detection by flow cytometry

Transfected HEK293T cells were stained with Fixable Viability Dye (eBioscience). After fixation (Fixation Buffer, Biolegend), cells were permeabilised (Perm Buffer, eBioscience) and stained with a monoclonal antibody that recognizes SARS-CoV-2 S1 (SinoBiological). Cells were acquired on a FACSCanto II flow cytometer (BD Biosciences) using BD FACSDiva software version 8.0.1 and analysed by FlowJo software version 10.6.2 (FlowJo LLC, BD Biosciences).

### Localization of expressed vaccine antigens by immunofluorescence

Transfected HEK293T cells were fixed in 4% paraformaldehyde (PFA) and permeabilised in PBS/0.2% Triton X-100. Free binding sites were blocked and cells incubated with a rabbit monoclonal antibody that recognizes the SARS-CoV-2 S1 subunit (SinoBiological), an anti-rabbit IgG secondary antibody (Jackson ImmunoResearch), labelled lectin HPA (Thermo Fisher Scientific) and concanavalin A (Fisher Scientific GmbH). DNA was stained with Hoechst (Life Technologies). Images were acquired with a Leica SP8 confocal microscope.

### SARS-CoV-2 RBD-foldon and P2 S expression and purification

To express the RBD-foldon encoded by BNT162b1 for ACE2 binding analysis and electron cryomicroscopy, DNA corresponding to the RNA coding sequence was cloned into the pMCG1309 vector. A plasmid encoding amino acids 1–615 of human ACE2 with C-terminal His-10 and Avi tags was generated for transient expression of the ACE2 peptidase domain (ACE2 PD) in Expi293F cells. The ACE2/B^0^AT1 complex was produced by co-expression of two plasmids in Expi293F cells, one of them encoding ACE2 amino acids 1–17 followed by haemagglutinin and Strep II tags and ACE2 amino acids 18–805, and the other containing a methionine followed by a FLAG tag and amino acids 2–634 of human B^0^AT1. Secreted ACE2 PD was isolated from conditioned cell culture medium using Nickel Excel resin (GE Healthcare) followed by gel filtration chromatography on a Superdex200 10/30 column (GE Healthcare) in PBS. Approximately 5 mg of purified ACE2 PD was covalently attached per 1 mL of 4% beaded agarose by amine coupling using AminoLink Plus resin (Thermo Fisher Scientific).

The RBD-trimer was purified from conditioned medium by affinity capture with the ACE2 PD crosslinked agarose and was eluted from the resin with 3 M MgCl2. Following dialysis, the protein was concentrated and purified by gel filtration using a Superdex200 10/300 column in 4-(2-hydroxyethyl)-1-piperazineethanesulfonic acid (HEPES)-buffered saline (HBS) with 10% glycerol. Purification of the ACE2/B^0^AT1 complex was based on the procedure described previously^5^. To form the ACE2/B^0^AT1/RBD-trimer complex, ACE2/B^0^AT1 aliquots were combined with purified RBD-foldon diluted in size exclusion chromatography buffer (25 mM Tris pH 8.0, 150 mM NaCl, 0.02% glyco diosgenin) for a 3:1 molar ratio of RBD-trimers to ACE2 protomers. After incubation at 4 °C for 30 minutes, the sample was concentrated and resolved on a Superose 6 Increase 10/300 GL column. Peak fractions containing the complex were pooled and concentrated.

To express SARS-CoV-2 P2 S encoded by BNT162b2 for characterisation by size exclusion chromatography, ACE2-PD binding, monoclonal antibody binding, and electron cryomicroscopy, a gene encoding the full length of SARS-CoV-2 (GenBank: MN908947) with two prolines substituted at residues 986 and 987 (K986P and V987P) followed with a C-terminal HRV3C protease site and a TwinStrep tag was cloned into a modified pcDNA3.1(+) vector with the CAG promoter. The TwinStrep-tagged P2 S was expressed in Expi293F cells.

Purification of the recombinant protein was based on a procedure described previously, with minor modifications^6^. Upon cell lysis, P2 S was solubilised in 1% NP-40 detergent. The TwinStrep-tagged protein was then captured with StrepTactin Sepharose HP resin in 0.5% NP-40. P2 S was further purified by size-exclusion chromatography and eluted as three distinct peaks in 0.02 % NP-40 as previously reported^6^. (Chromatogram not shown.) A peak that consists of intact P2 S migrating at around 150 kDa, as well as dissociated S1 and S2 subunits (which co-migrate at just above 75 kDa), was used in the structural characterisation. Spontaneous dissociation of the S1 and S2 subunits occurs throughout the course of protein purification, starting at the point of detergent-mediated protein extraction, so that P2 S preparations also contain dissociated S1 and S2.

### Binding kinetics of the RBD-foldon trimer and P2 S to immobilised human ACE2 and a neutralizing monoclonal antibody by biolayer interferometry

Binding of purified RBD-foldon to the human ACE2 peptidase domain (ACE2 PD) and of NP-40 solubilised, purified P2 S to ACE2-PD and human neutralising monoclonal antibody B38^25^ was measured by biolayer interferometry at 25 °C on an Octet RED384 (FortéBio). RBD-foldon binding was measured in 10 mM HEPES pH 7.5, 150 mM NaCl and 1 mM ethylenediaminetetraacetic acid (EDTA). P2 S binding was measured in 25 mM Tris pH 7.5, 150 mM NaCl, 1 mM EDTA and 0.02% NP-40. Avi-tagged human ACE2 PD was immobilised on streptavidin-coated sensors; Avi-tagged B38 antibody was immobilised on protein G-coated sensors. For a RBD-foldon concentration series, binding data were collected for 600 seconds of association and 900 seconds of dissociation. For a P2 S concentration series, after initial baseline equilibration of 120 seconds, the sensors were dipped in a 10 µg/mL solution of Avi-tagged ACE2-PD or B38 mAb for 300 seconds to achieve capture levels of 1 nM using the threshold function. Then, after another 120 seconds of baseline, binding data were collected for 300 seconds of association and 600 seconds of dissociation.

Biolayer interferometry data were collected with Octet Data Acquisition software version 10.0.0.87 and processed using ForteBio Data Analysis software version 10.0. Data were reference subtracted and fit to a 1:1 binding model with R^2^ value greater than 0.96 for the RBD and 0.95 for P2 S to determine kinetics and affinity (P2 S) or avidity (RBD-foldon) of binding using Octet Data Analysis Software v10.0 (FortéBio). For the RBD-foldon, the dissociation rate of interaction (*k*d) with ACE2-PD was slower than the limit of measurement of the instrument, and the minimum binding avidity (*K*D) was estimated using an assumed dissociation rate *k*d of 1 × 10^-6^ s^-1^.

### Electron microscopy of negatively stained RBD-foldon trimers

Purified RBD-foldon in 4 μL was applied to a glow-discharged copper grid overlaid with formvar and amorphous carbon (Ted Pella). Negative staining was performed with Nano-W organotungstate stain (Nanoprobes) according to the manufacturer’s protocol. The sample imaged using an FEI TF-20 microscope operating at 200 kV, with a magnification of 62,000x and defocus of −2.5 μm. Micrographs were contrast transfer function (CTF)-corrected in RELION using CTFFIND-4.1^43^. A small manually picked dataset was used to generate 2D references for auto-picking. The resulting particle set was subjected to 2D classification in RELION 3.0.6^44^.

### Cryo-EM of the ACE2/B^0^AT1/RBD-trimer complex

Cryo-EM was performed using a Titan Krios operating at 300 keV equipped with a Gatan K2 Summit direct electron detector in super-resolution mode at a magnification of 165,000x, for a magnified pixel size of 0.435 Å at the specimen level.

Purified ACE2/B^0^AT1/RBD-trimer complex at 6 mg/mL in 4 μL was applied to gold Quantifoil R1.2/1.3 200 mesh grids glow discharged in residual air for 30 seconds at 20 mA using a Pelco Easiglow. The sample was blotted using a Vitrobot Mark IV for 5 seconds with a force of −3 before being plunged into liquid ethane cooled by liquid nitrogen. In total, 7,455 micrographs were collected from a single grid. Data were collected over a defocus range of −1.2 to −3.4 μm with a total electron dose of 52.06 e^-^/Å^2^ fractionated into 40 frames over a 6-second exposure for 1.30 e^-^/Å^2^/frame. Initial motion correction was performed in Warp^45^, during which super-resolution data were binned to give a pixel size of 0.87 Å. Corrected micrographs were imported into RELION 3.1-beta^44^ for CTF estimation with CTFFIND-4.1^43^.

Particles were picked using the LaPlacian-of-Gaussian particle picking algorithm as implemented in RELION and extracted with a box size of 450 pixels. References obtained by 2D classification were used for a second round of reference-based auto-picking, yielding a dataset of 715,356 particles. Two of the three RBDs of each particle (the two not constrained by binding to ACE2/B^0^AT1) exhibited diffuse density in 2D classification that reflected high particle flexibility, consistent with the conformational flexibility of RBD trimers observed by negative stain EM (Fig. 1c, d). This flexibility precluded the inclusion of all three RBDs in the final structural solution. Particle heterogeneity was filtered out with 2D and 3D classification with a mask size of 280 Å to filter out the diffuse density of the two non-ACE2-bound RBD copies in each RBD-trimer, yielding a set of 87,487 particles, which refined to 3.73 Å with C2 symmetry. Refinement after subtraction of micelle and B^0^AT1 density from the particles yielded an improved map of 3.24 Å. The atomic model from PDB ID 6M17^5^ was rigid-body fitted into the 3.24 Å density and then flexibly fitted to the density using real-space refinement in Phenix^46^ alternating with manual building in Coot^47^. The microscope was operated for image acquisition using SerialEM software version 3.8.0 beta^48^. Validation of this model is shown in Supplementary Fig. 2. Data collection, 3D reconstruction and model refinement statistics are listed in Extended Data Table 1.

### Cryo-EM of P2 S

For TwinStrep-tagged P2 S, 4 μL purified protein at 0.5 mg/mL were applied to gold Quantifoil R1.2/1.3 300 mesh grids freshly overlaid with graphene oxide. The sample was blotted using a Vitrobot Mark IV for 4 seconds with a force of −2 before being plunged into liquid ethane cooled by liquid nitrogen. 27,701 micrographs were collected from two identically prepared grids. Data were collected from each grid over a defocus range of −1.2 to −3.4 μm with a total electron dose of 50.32 and 50.12 e^-^/Å^2^, respectively, fractionated into 40 frames over a 6-second exposure for 1.26 and 1.25 e^-^/Å^2^/frame. On-the-fly motion correction, CTF estimation, and particle picking and extraction with a box size of 450 pixels were performed in Warp^45^, during which super-resolution data were binned to give a pixel size of 0.87 Å. A total of 1,119,906 particles were extracted. All subsequent processing was performed in RELION 3.1-beta^44^. Particle heterogeneity was filtered out with 2D and 3D classification, yielding a set of 73,393 particles, which refined to 3.6 Å with C3 symmetry. 3D classification of this dataset without particle alignment separated out one class with a single RBD up, representing 15,098 particles. The remaining 58,295 particles, in the three RBD ‘down’ conformation, were refined to give a final model at 3.29 Å. The atomic model from PDB ID 6XR8^6^ was rigid-body fitted into the map density, then flexibly fitted to the density using real-space refinement in Phenix^46^ alternating with manual building in Coot^47^. The cryo-EM model validation is provided in Extended Data Fig. 2, the full cryo-EM data processing workflow, and the model refinement statistics in Extended Data Table. 1.

### Immunisation

#### Mice

Female BALB/c mice (Janvier; 8-12 weeks) were randomly allocated to groups. BNT162b1 and BNT162b2 were diluted in PBS with 300 mM sucrose (Fig. 2 and Fig. 3b, d for BNT162b2, and Extended Data Fig. 3) or 0.9% NaCl placebo control (Fig. 3a, c and Fig. 3b, d for BNT162b1, and Extended Data Fig. 4) and injected IM into the gastrocnemius muscle at a volume of 20 µL under isoflurane anaesthesia.

#### Rhesus macaques (Macaca mulatta)

Male rhesus macaques (2–4 years old) were randomly assigned to receive BNT162b1 or BNT162b2 on Days 0 and 21 or saline control on Days 0 and 21 or 35. Vaccine was administered in 0.5 mL by IM injection in the left quadriceps muscle. Animals were anesthetised with ketamine HCl (10 mg/kg; IM) during immunisation and were monitored for adequate sedation.

### Phlebotomy and tissue preparation

#### Mice

Peripheral blood was collected from the retro-orbital venous plexus under isoflurane anaesthesia or *vena facialis* without anaesthesia. For flow cytometry, blood was heparinised. For serum generation, blood was centrifuged for 5 min at 16,000 x g, and the serum was immediately used for downstream assays or stored at −20 °C. Spleen single-cell suspensions were prepared in PBS by mashing tissue against the surface of a 70 µm cell strainer (BD Falcon). Erythrocytes were removed by hypotonic lysis. Popliteal, inguinal and iliac lymph nodes were pooled, cut into pieces, digested with collagenase D (1 mg/mL; Roche) and passed through cell strainers.

#### Rhesus macaques (Macaca mulatta)

Serum was obtained before, 6 hours after, and 1, 14, 21, 28, 35 and 42 days after immunisation with BNT162b1, BNT162b2, or saline (Extended Data Table 2). For BNT162b2 and challenge cohort 3 controls, serum was also obtained on Day 56, and PBMCs were obtained before immunisation and on Days 7, 28, and 42, except that PBMCs were not obtained from the challenge cohort 3 control animals on Day 28. Blood for serum and PBMCs was collected in compliance with animal protocol 2017-8725-023 approved by the NIRC Institutional Animal Care and Use Committee. Animals were anesthetised with ketamine HCl (10 mg/kg; IM) during blood collection and were monitored for adequate sedation.

### Analysis of S1- and RBD-specific serum IgG

#### Mice

MaxiSorp plates (Thermo Fisher Scientific) were coated with recombinant S1 or RBD (1 µg/mL) in sodium carbonate buffer, and serum-derived, bound IgG was detected using a horseradish peroxidase (HRP)-conjugated secondary antibody and tetramethylbenzidine (TMB) substrate (Biotrend). Data collection was performed using a BioTek Epoch reader and Gen5 software version 3.0.9. For concentration analysis, an IgG mouse isotype control was used in parallel in a serial dilution, and the sample signals were correlated to a standard curve of the isotype control.

#### Rhesus macaques (Macaca mulatta), humans

Recombinant SARS-CoV-2 S1 containing a C-terminal Avitag™ (Acro Biosystems) was bound to streptavidin-coated Luminex microspheres. Bound rhesus macaque or human anti-S1 antibodies present in the serum were detected with a fluorescently labelled goat anti-human polyclonal secondary antibody (Jackson ImmunoResearch). Data were captured as median fluorescent intensities (MFIs) using a Bioplex200 system (Bio-Rad) and converted to U/mL antibody concentrations using a reference standard consisting of 5 pooled human COVID-19 convalescent serum samples (obtained >14 days PCR diagnosis, from the panel described above), diluted in antibody depleted human serum with arbitrary assigned concentrations of 100 U/mL and accounting for the serum dilution factor.

### Surface plasmon resonance spectroscopy of polyclonal mouse immune sera

Binding kinetics of murine S1- and RBD-specific serum IgG to recombinant S1 and RBD was determined using a Biacore T200 device (Cytiva) with 10 mM Hepes, 150 mM NaCl, 3 mM EDTA, 0.05% v/v surfactant P20 (HBS-EP running buffer, BR100669, Cytiva) at 25 °C. Carboxyl groups on the CM5 sensor chip matrix were activated with a mixture of 1-ethyl-3-(3-dimethylaminopropyl) carbodiimidehydrochloride (EDC) and N-hydroxysuccinimide (NHS) to form active esters for the reaction with amine groups. Anti-mouse-Fc-antibody (Jackson ImmunoResearch) was diluted in 10 mM sodium acetate buffer pH 5 (30 µg/mL) for covalent coupling to immobilisation level of ∼10,000 response units (RU). Free N-hydroxysuccinimide esters on the sensor surface were deactivated with ethanolamine.

Mouse serum was diluted 1:50 in HBS-EP buffer and applied at 10 µL/min for 30 seconds to the active flow cell for capture by immobilised antibody, while the reference flow cell was treated with buffer. Binding analysis of captured murine IgG antibodies to S1-His or RBD-His (Sino Biological Inc.) was performed using a multi-cycle kinetic method with concentrations ranging from 25 to 400 nM or 1.5625 to 50 nM, respectively. An association period of 180 seconds was followed by a dissociation period of 600 seconds with a constant flow rate of 40 μL/min and a final regeneration step. Binding kinetics were calculated using a global kinetic fit model (1:1 Langmuir, Biacore T200 Evaluation Software Version 3.1, Cytiva).

### VSV-SARS-CoV-2 S pseudovirus entry inhibition assay by serum IgG in mice

A recombinant replication-deficient vesicular stomatitis virus (VSV) vector that encodes green fluorescent protein (GFP) instead of VSV-G (VSVΔG-GFP) was pseudotyped with SARS-CoV-2 S according to published pseudotyping protocols^49, 50^. In brief, HEK293T/17 monolayers transfected to express SARS-CoV-2 S truncated of the C-terminal cytoplasmic 19 amino acids (SARS-CoV-2-S-CΔ19) were inoculated with VSVΔG-GFP vector (rescued from pVSVΔG-GFP plasmid expression vector; Kerafast Inc.). After incubation for 1 h at 37 °C, the inoculums was removed, and cells were washed with PBS before medium supplemented with anti-VSV-G antibody (clone 8G5F11, Kerafast Inc.) was added to neutralise residual input virus. VSV/SARS-CoV-2 pseudovirus-containing medium was harvested 20 h after inoculation, 0.2 µm filtered and stored at −80 °C.

Vero-76 cells were seeded in 96-well plates. Serial dilutions of mouse serum samples were prepared and pre-incubated for 10 min at room temperature with VSV/SARS-CoV-2 pseudovirus suspension (4.8 × 10^3^ infectious units [IU]/mL) before transferring the mix to Vero-76 cells. Inoculated Vero-76 cells were incubated for 20 h at 37 °C. Plates were placed in an IncuCyte Live Cell Analysis system (Sartorius) and incubated for 30 min prior to the analysis (IncuCyte 2019B Rev2 software). Whole well scanning for brightfield and GFP fluorescence was performed using a 4× objective. The 50% pseudovirus neutralisation titre (pVNT50) was reported as the reciprocal of the highest dilution of serum still yielding a 50% reduction in GFP-positive infected cell number per well compared to the mean of the no serum pseudovirus positive control. Each serum sample dilution was tested in duplicates.

### IFNγ and IL-4 ELISpot

#### Mice

ELISpot assays were performed with mouse IFNγ ELISpot^PLUS^ kits according to the manufacturer’s instructions (Mabtech). A total of 5 × 10^5^ splenocytes was *ex vivo* restimulated with the full-length S peptide mix (0.1 µg/mL final concentration per peptide) or controls (gp70-AH1 [SPSYVYHQF]^38^, 4 µg/mL; concanavalin A [ConA], 2 µg/mL [Sigma]). Streptavidin-alkaline phosphatase (ALP) and 5-bromo-4-chloro-3′-indolyl phosphate (BCIP)/nitro blue tetrazolium (NBT)-plus substrate were added, and spots counted using an ELISpot plate reader (ImmunoSpot® S6 Core Analyzer [CTL]). Spot numbers were evaluated using ImmunoCapture Image Acquisition Software V7.0 and ImmunoSpot 7.0.17.0 Professional. Spot counts denoted too numerous to count by the software were set to 1,500. For T-cell subtyping, CD8^+^ T cells and CD4^+^ T cells were isolated from splenocyte suspensions using MACS MicroBeads (CD8a [Ly-2] and CD4 [L3T4] [Miltenyi Biotec]) according to the manufacturer’s instructions. CD8^+^ or CD4^+^ T cells (1 × 10^5^) were subsequently re-stimulated with 5 × 10^4^ syngeneic bone marrow-derived dendritic cells loaded with full-length S peptide mix (0.1 µg/mL final concentration per peptide), or cell culture medium as control. Purity of isolated T-cell subsets was determined by flow cytometry to calculate spot counts per 1 × 10^5^ CD8^+^ or CD4^+^ T cells.

#### Rhesus macaques (Macaca mulatta)

Rhesus macaque PBMCs were tested with commercially available NHP IFNγ and IL-4 ELISpot assay kits (Mabtech). Cryopreserved rhesus macaque PBMCs were thawed in pre-warmed AIM-V media (Thermo Fisher Scientific) with Benzonase (EMD Millipore). For IFNγ ELISpot, 1.0 x 10^5^ PBMCs and for IL-4 ELISpot, 2.5 x 10^5^ PBMCs were stimulated *ex vivo* with 1 μg/mL of the full-length S overlapping peptide mix. Tests were performed in triplicate wells and medium containing dimethyl sulphoxide (media-DMSO), a CMV peptide pool and phytohemagglutinin (PHA; Sigma) were included as controls. After 24 h for IFNγ and 48 h for IL-4, streptavidin-HRP and 3-amino-9-ethylcarbazole (AEC) substrate (BD Bioscience) were added and spots counted using a CTL ImmunoSpot S6 Universal Analyzer (CTL). Results shown are background (Medium-DMSO) subtracted and normalised to SFC/10^6^ PBMCs.

### Cell-mediated immunity by flow cytometry

#### Mice

For T-cell analysis in peripheral blood, erythrocytes from 50 µL freshly drawn blood were lysed (ammonium-chloride-potassium lysing buffer [Gibco]), and cells were stained with Fixable Viability Dye (eBioscience) and primary antibodies in the presence of Fc block in flow buffer (Dulbecco’s phosphate-buffered saline [Gibco] supplemented with 2% fetal calf serum (FCS), 2 mM ethylenediaminetetraacetic acid [both Sigma] and 0.01% sodium azide [Morphisto]). After staining with secondary biotin-coupled antibodies in flow buffer, cells were stained extracellularly against surface markers with directly labelled antibodies and streptavidin in Brilliant Stain Buffer Plus (BD Bioscience) diluted in flow buffer. Cells were washed with 2% RotiHistofix (Carl Roth), fixed (Fix/Perm Buffer, FoxP3/Transcription Factor Staining Buffer Set [eBioscience]) and permeabilised (Perm Buffer, FoxP3/Transcription Factor Staining Buffer Set [eBioscience]) overnight. Permeabilised cells were intracellularly treated with Fc block and stained with antibodies against transcription factors in Perm Buffer.

For T-cell analysis in lymphoid tissues, 1 × 10^6^ lymph node cells (for BNT162b1) or 1.5 × 10^6^ lymph node cells (for BNT162b2) and 4 × 10^6^ spleen cells were stained for viability and extracellular antigens with directly labelled antibodies. Fixation, permeabilisation and intracellular staining was performed as described for blood T-cell staining.

For B-cell subtyping in lymphoid tissues, 2.5 × 10^5^ lymph node and 1 × 10^6^ spleen cells were treated with Fc block, stained for viability and extracellular antigens as described for blood T-cell staining and fixed with 2% RotiHistofix overnight.

For intracellular cytokine staining of T cells from BNT162b1-immunised mice, 1 x 10^6^ lymph node and 4 x 10^6^ spleen cells were *ex vivo* restimulated with 0.2 µg/mL final concentration per peptide of full-length S peptide mix. For intracellular cytokine staining of T cells from mice immunised with BNT162b2, 4 x 10^6^ spleen cells were *ex vivo* restimulated with 0.5 µg/mL final concentration per peptide of full-length S peptide mix or cell culture medium (no peptide) as control. The cells were restimulated for 5 hours in the presence of GolgiStop and GolgiPlug (both BD Bioscience) for 5 hours. Cells were stained for viability and extracellular antigens as described for lymphoid T-cell staining. Cells were fixed with 2% RotiHistofix and permeabilised overnight. Intracellular staining was performed as described for blood T-cell staining.

Mouse cells were acquired on a BD Symphony A3 or BD Celesta (B-cell subtyping) flow cytometer (BD Bioscience) using BD FACSDiva software version 9.1 or 8.0.1.1, respectively, and analysed with FlowJo 10.6 (FlowJo LLC, BD Biosciences).

#### Rhesus macaques (Macaca mulatta)

For intracellular cytokine staining in T cells, 1.5 x 10^6^ PBMCs were stimulated with the full-length S peptide mix at 1 μg/mL (concentration of all peptides, combined), Staphyloccocus enterotoxin B (SEB; 2 μg/mL) as positive control, or 0.2% DMSO as negative control. GolgiStop and GolgiPlug (both BD Bioscience) were added. Following 37 °C incubation for 12 to 16 h, cells were stained for viability and extracellular antigens after blocking Fc binding sites with directly labelled antibodies. Cells were fixed, permeabilised with BDCytoFix/CytoPerm solution (BD Bioscience), and intracellular staining was performed in the permeabilisation buffer for 30 min at room temperature. Cells were washed, resuspended in 2% FBS/PBS buffer and acquired on an LSR Fortessa. Data were analysed by FlowJo 10.4.1 (FlowJo LLC, BD Biosciences). Results shown are background (media-DMSO) subtracted.

### Cytokine profiling in mice by bead-based immunoassay

Mouse splenocytes were re-stimulated for 48 h with full-length S peptide mix (0.1 µg/mL final concentration per peptide) or cell culture medium (no peptide) as control. Concentrations of IFNγ, IL-2, IL-4, IL-5 and (for splenocytes from BNT162b2-immunised mice) IL-13 in supernatants were determined using a bead-based, 11-plex TH1/TH2 mouse ProcartaPlex multiplex immunoassay (Thermo Fisher Scientific) according to the manufacturer’s instructions. Fluorescence was measured with a Bioplex200 system (Bio-Rad) and analysed with ProcartaPlex Analyst 1.0 software (Thermo Fisher Scientific). Values below the lower limit of quantification (LLOQ) were set to zero.

### SARS-CoV-2 neutralisation by rhesus macaque (*Macaca mulatta*) sera

The SARS-CoV-2 neutralisation assay used a previously described strain of SARS-CoV-2 (USA_WA1/2020) that had been rescued by reverse genetics and engineered by the insertion of an mNeonGreen (mNG) gene into open reading frame 7 of the viral genome^27^. This reporter virus generates similar plaque morphologies and indistinguishable growth curves from wild-type virus. Viral master stocks were grown in Vero E6 cells as previously described^51^. When testing human convalescent serum specimens, the fluorescent neutralisation assay produced comparable results to the conventional plaque reduction neutralisation assay. Serial dilutions of heat-inactivated sera were incubated with the reporter virus (2 x 10^4^ plaque forming units [PFU] per well) to yield an approximately 10-30% infection rate of the Vero CCL81 monolayer for 1 h at 37 °C before inoculating Vero CCL81 cell monolayers (targeted to have 8,000 to 15,000 cells in the central field of each well at the time of seeding, one day before infection) in 96-well plates to allow accurate quantification of infected cells. Cell counts were enumerated by nuclear stain (Hoechst 33342), and fluorescent virus-infected foci were detected 16-24 hours after inoculation with a Cytation 7 Cell Imaging Multi-Mode Reader (BioTek) with Gen5 Image Prime version 3.09. Titres were calculated in GraphPad Prism version 8.4.2 by generating a 4-parameter (4PL) logistical fit of the percent neutralisation at each serial serum dilution. The 50% neutralisation titre (VNT50) was reported as the interpolated reciprocal of the dilution yielding a 50% reduction in fluorescent viral foci.

### SARS-CoV-2 challenge of rhesus macaques (*Macaca mulatta*)

The SARS-CoV-2 inoculum was obtained from a stock of 2.1 × 10^6^ PFU/mL previously prepared at Texas Biomedical Research Institute (San Antonio, TX), aliquoted into single use vials, and stored at −70 °C. The working virus stock was generated from two passages of the SARS-CoV-2 USA-WA1/2020 isolate (a 4^th^ passage seed stock purchased from BEI Resources; NR-52281) in Vero E6 cells. The virus was confirmed to be SARS-CoV-2 by deep sequencing that demonstrated identity to a published SARS-CoV-2 sequence (GenBank accession number MN985325.1).

BNT162b1-immunised (*n*=6), BNT162b2-immunised (*n*=6), and age-matched saline-immunised (*n*=9) male rhesus macaques (control) were challenged with 1.05 × 10^6^ plaque forming units of SARS-CoV-2 USA-WA1/2020 isolate, split equally between the intranasal (IN; 0.25 mL) and intratracheal (IT; 0.25 mL) routes as previously described^28^. Sentinel age- and sex-matched animals (*n*=6) were mock challenged with DMEM supplemented with 10% FCS IN (0.25 mL) and IT (0.25 mL). The macaques were challenged or mock challenged at the times relative to immunisation indicated in Extended Data Fig. 6 and Extended Data Table 2.

Twelve to nineteen days prior to challenge, animals were moved from the NIRC, in New Iberia, LA, where they had been immunised, to the animal biosafety level 3 facility at SNPRC (in San Antonio, TX). Animals were monitored regularly by a board-certified veterinary clinician for rectal body temperature, weight and physical examination. Specimen collection was performed under tiletamine zolazepam (Telazol) anaesthesia as described^28^. Bronchoalveolar lavage (BAL), nasal, OP and rectal swab collection, X-ray and CT examinations and necropsy were performed at the times indicated in Extended Data Figure 6 and Extended Data Table 2. The 3 control animals in challenge cohort 3 and 3 sentinel animals were not necropsied to allow their subsequent re-challenge (control) or challenge (sentinel). BAL was performed by instilling 20 mL of saline 4 times. These washings were pooled, aliquoted and stored frozen at −70 °C.

### SARS-CoV-2 viral RNA quantification by reverse-transcription quantitative polymerase chain reaction

To detect and quantify SARS-CoV-2 in NHP, viral RNA was extracted from BAL fluid and from nasal, OP, and rectal swabs as previously described^52–54^ and tested by RT-qPCR as previously described^28^. Briefly, 10 µg yeast tRNA and 1 × 10^3^ PFU of MS2 phage (*Escherichia coli* bacteriophage MS2, ATCC) were added to each thawed sample, and RNA extraction performed using the NucleoMag Pathogen kit (Macherey-Nagel). The SARS-CoV-2 RT-qPCR was performed on extracted RNA using a CDC-developed 2019-nCoV N1 assay on a QuantStudio 3 instrument (Applied Biosystems). The cut-off for positivity (limit of detection, LOD) was established at 10 gene equivalents (GE) per reaction (800 GE/mL). Samples were tested in duplicate. One BAL specimen from the challenge cohort 2 control group obtained on Day 6 after challenge and one nasal swab from the BNT162b1-immunised group obtained on Day 1 after challenge had, on repeated measurements, viral RNA levels on either side of the LLOD. These specimens were categorised as indeterminate and excluded from the graphs and the analysis.

### Radiology

Thoracic radiographs and computed tomography (CT) scans were performed under anesthesia as previously described^28^. For radiographic imaging, 3-view thoracic radiographs (ventrodorsal, right and left lateral) were obtained at the times relative to challenge indicated in Extended Data Table 2. The animals were anesthetized using Telazol (2-6 mg/kg) and maintained by inhaled isoflurane delivered through a Hallowell 2002 ventilator anesthesia system (Hallowell, Pittsfield, MA). Animals were intubated to perform end inspiratory breath-hold using a remote breath-hold switch. Lung field CT images were acquired using Multiscan LFER150 PET/CT (MEDISO Inc., Budapest, Hungary) scanner. Image analysis was performed using 3D ROI tools available in Vivoquant (Invicro, Boston, MA). Images were interpreted by a board-certified veterinary radiologist blinded to treatment groups. Scores were assigned to a total of 7 lung regions on a severity scale of 0-3 per region, with a maximum severity score of 21. Pulmonary lesions evident prior to challenge, or those which could not be unequivocally attributed to the viral challenge (such as atelectasis secondary to recumbency and anesthesia) received a score of “0”.

### Histopathology

Lung histopathology is reported on necropsies performed on 2-4 year old male rhesus macaques at the times after challenge indicated in Extended Data Figure 6 and Extended Data Table 2. Necropsy, tissue processing, and histology were performed by SNPRC in San Antonio, TX. Samples were fixed in 10% neutral buffered formalin and processed routinely into paraffin blocks. Tissue blocks were sectioned to 5 µm and stained with hematoxylin and eosin. Microscopic evaluation of 7 lung tissue sections per animal (1 sample of each lobe on L & R) was performed blindly by SNPRC and Pfizer pathologists. Lungs were evaluated using a semi-quantitative scoring system with inclusion of cell types and/or distribution as appropriate. Inflammation score was based on area of tissue in section involved: 0 = normal; 1=<10%; 2=11-30%; 3=30-60%; 4= 60-80%; 5=>80%. Each lobe received an individual score, and the final score for each animal was reported as the mean of the individual scores. The pathologists were unblinded to the group assignments after agreement on diagnoses. As indicated in Extended Data Fig. 6 and Extended Data Table 2, the BNT162b1-immunised and control macaques were challenged and necropsied in parallel (challenge cohorts 1 and 2), and the BNT162b2-immunised rhesus macaques were immunised and challenged subsequently (challenge cohort 3).

### Statistics and reproducibility

No statistical methods were used to predetermine group and samples sizes (*n*). All experiments were performed once. P-values reported for RT-qPCR analysis were determined by nonparametric analysis (Friedman’s test) based on the ranking of viral RNA shedding data within each day. PROC RANK and PROC GLM from SAS® 9.4 were used to calculate the p-values. All available post-challenge BAL fluid and nasal, OP, and rectal swab samples from the necropsied animals and all available post-challenge samples through Day 10 from the animals not necropsied were included in the analysis. Indeterminate results were excluded from this analysis. All remaining analyses were two-tailed and carried out using GraphPad Prism 8.4.

### Data availability

The data that support the findings of this study are available from the corresponding author upon reasonable request.

## Acknowledgements

We thank T. Garretson and D. Cooper for advice on and M. Cutler for coordination of NHP serology studies. We thank R. F. Sommese and K. F. Fennell for technical assistance for molecular cloning and cell-based binding. Valuable support and assistance M. Dvorak, M. Drude, F. Zehner, T. Lapin, B. Ludloff, S. Hinz, F. Bayer, J. Scholz, A.L. Ernst, T. Sticker and S. Wittig resulted in a rapid availability of oligonucleotides and DNA templates. E. Boehm, K. Goebel, R. Frieling, C. Berger, S. Koch, T. Wachtel, J. Leilich, M. Mechler, R. Wysocki, M. Le Gall, A. Czech and S. Klenk carried out RNA production and analysis. Without their commitment during this pandemic situation, this vaccine candidate could not have been transferred to non-clinical studies in light speed. B. Weber, J. Vogt, S. Krapp, K. Zwadlo, J. Mottl, J. Mühl and P. Windecker supported the mouse studies and serological analysis with excellent technical assistance. We thank radiologists A. K. Voges and E. Clemmons for interpreting radiographs for the nonhuman primate study and S. Ganatra for radiology services. We thank E. Romero for veterinary services and K.A. Soileau and the staff of the New Iberia Research Centre for non-human primate care. We thank E.J. Dick for veterinary pathology. We thank Polymun Scientific for excellent formulation services as well as Acuitas Therapeutics for fruitful discussions. We thank S. Wigge and C. Lindemann for scientific writing support and manuscript review.

## Author Contributions

U.S. conceived and conceptualised the work and strategy. S.H., S.C.D., A.A.H.S., C.G., R.d.l.C.G.G., and M.C.G. designed primers, performed oligosynthesis, cloned constructs and performed protein expression experiments. T.Z., S.F., J.S. and A.N.K. developed, planned, performed and supervised RNA synthesis and analysis. E.H.M. purified P2 S. N.L.N. purified RBD-trimer and ACE2 PD. J.A.L. developed ACE2/B^0^AT1/RBD-trimer formation and purified the complex. P.V.S. developed and performed biolayer interferometry experiments. J.A.L. and S.H. performed electron microscopy and solved the structure of the complex. Y.C. supervised the structural and biophysical characterisation and analysed the structures. A.M. and B.G.L. performed surface plasmon resonance spectroscopy. A.G., S.A.K, S.S., T.H., L.F. and F.V. planned, performed and analysed *in vitro* studies. F.B., T.K., C.R. managed formulation strategy. A.B.V., M.V., L.M.K., K.C.W. designed mouse studies, analysed and interpreted data. A.P., S.E., D.P. and G.S. performed and analysed the S1- and RBD-binding IgG assays. M.G. designed and optimized MS2-SARS-nCoV-2-N1 RT-qPCR assay. M.G., R.C., Jr., and K.J.A. performed and analysed viral RT-qPCR data. A.M., B.S. and A.W. performed and analysed pVNT, C.F.-G. and P.-Y.S performed and analysed VNT assays. D.E., D.S., B.J., Y.F, H.J. performed *in vivo* studies and ELISpot assays. A.B.V., K.C.W. J.L., M.S.M. A.O.-S., and M.V. planned, analysed and interpreted ELISpot assays. L.M.K., J.L., D.E., Y.F., H.J., A.P.H. M.S.M. and P.A. planned, performed and analysed flow cytometry assays. A.B.V., L.M.K., Y.F. and H.J. planned, performed, analysed and interpreted cytokine release assays. M.R.G. read and interpreted radiographs and CT scan. O.G. and S.C. read and interpreted histopathology specimens. R.S.S. and S.C. interpreted histopathology data. I.K., K.A.S., K.T., C.Y.T., M.G., D.K. and P.R.D. designed NHP studies, analysed and interpreted data. K.T., M.P., I.L.S. and W.K. oversaw NHP immunogenicity and serology testing. S.H.-U. and K.B. provided veterinary services for NHPs. J.A.F., J.C., T.C. and J.O. managed the NHP colony. U.S., Ö.T., P.R.D, L.M.K., A.M., M.V. contributed to synthesis and integrated interpretation of obtained data. A.B.V., I.K., Y.C., A.M., M.V, L.M.K., C.T., K.A.S., Ö.T., P.R.D, K.U.J. and U.S. wrote the manuscript. All authors supported the review of the manuscript.

## Additional Information

Supplementary Information is available for this study.

Correspondence and requests for materials should be addressed to Ugur Sahin.

---

**Extended Data Figure 1.**
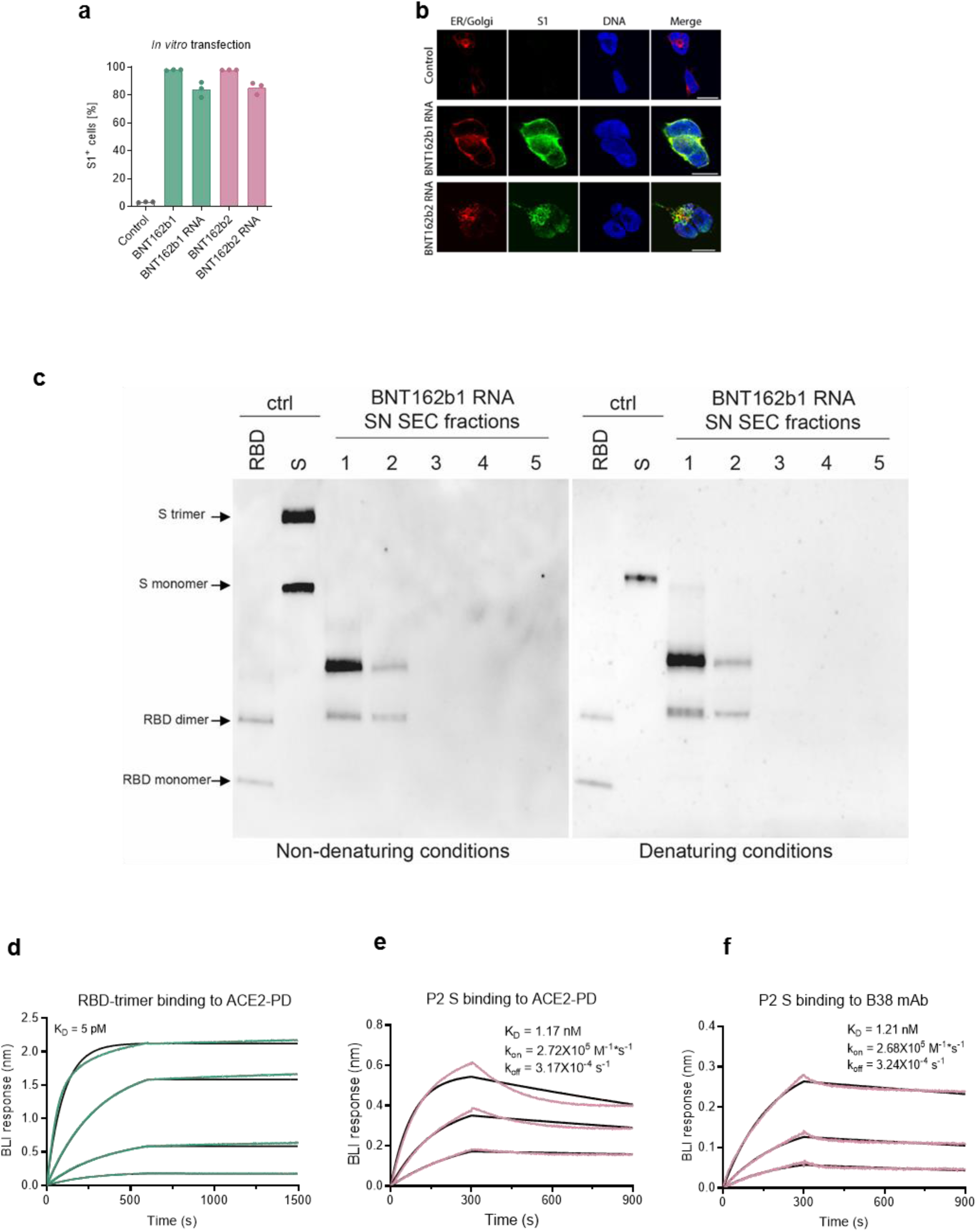
Vaccine antigen expression and receptor affinity. **a**, Detection of BNT162b1-encoded RBD-foldon and BNT162b2-encoded P2 S in HEK293T cells by S1-specific antibody staining and flow cytometry. HEK293T cells analysed by flow cytometry were incubated with: no RNA (control), BNT162b RNAs formulated as LNPs (BNT162b1, BNT162b2) or BNT162b RNAs mixed with a transfection reagent (BNT162b1 RNA, BNT162b2 RNA). **b**, Localisation of BNT162b1 RNA-encoded RBD-foldon or BNT162b2 RNA-encoded P2 S in HEK293T cells transfected as in panel a, determined by immunofluorescence staining. Endoplasmic reticulum and Golgi (ER/Golgi, red), S1 (green) and DNA (blue). Scale bar: 10 µm. **c**, Western blot of denatured and non-denatured samples of size exclusion chromatography (SEC) fractions (chromatogram in Supplementary Fig. 1) of concentrated medium from HEK293T cells transfected with BNT162b1 RNA. The RBD-foldon was detected with a rabbit monoclonal antibody against the S1 fragment of SARS-CoV-2 S. Protein controls (ctrl): purified, recombinant RBD and S. **d,** Biolayer interferometry sensorgram demonstrating the binding kinetics of the purified RBD-foldon trimer, expressed from DNA, to immobilised human ACE2-PD. **e**,**f** Biolayer inferometry sensorgrams showing binding of a DNA-expressed P2 S preparation from a size exclusion chromatography peak (not shown) that contains intact P2 S and dissociated S1 and S2 to immobilised (**e**) human ACE2-PD and (**f**) B38 monoclonal antibody. Binding data are in colour; 1:1 binding models fit to the data are in black.

**Extended Data Figure 2.**
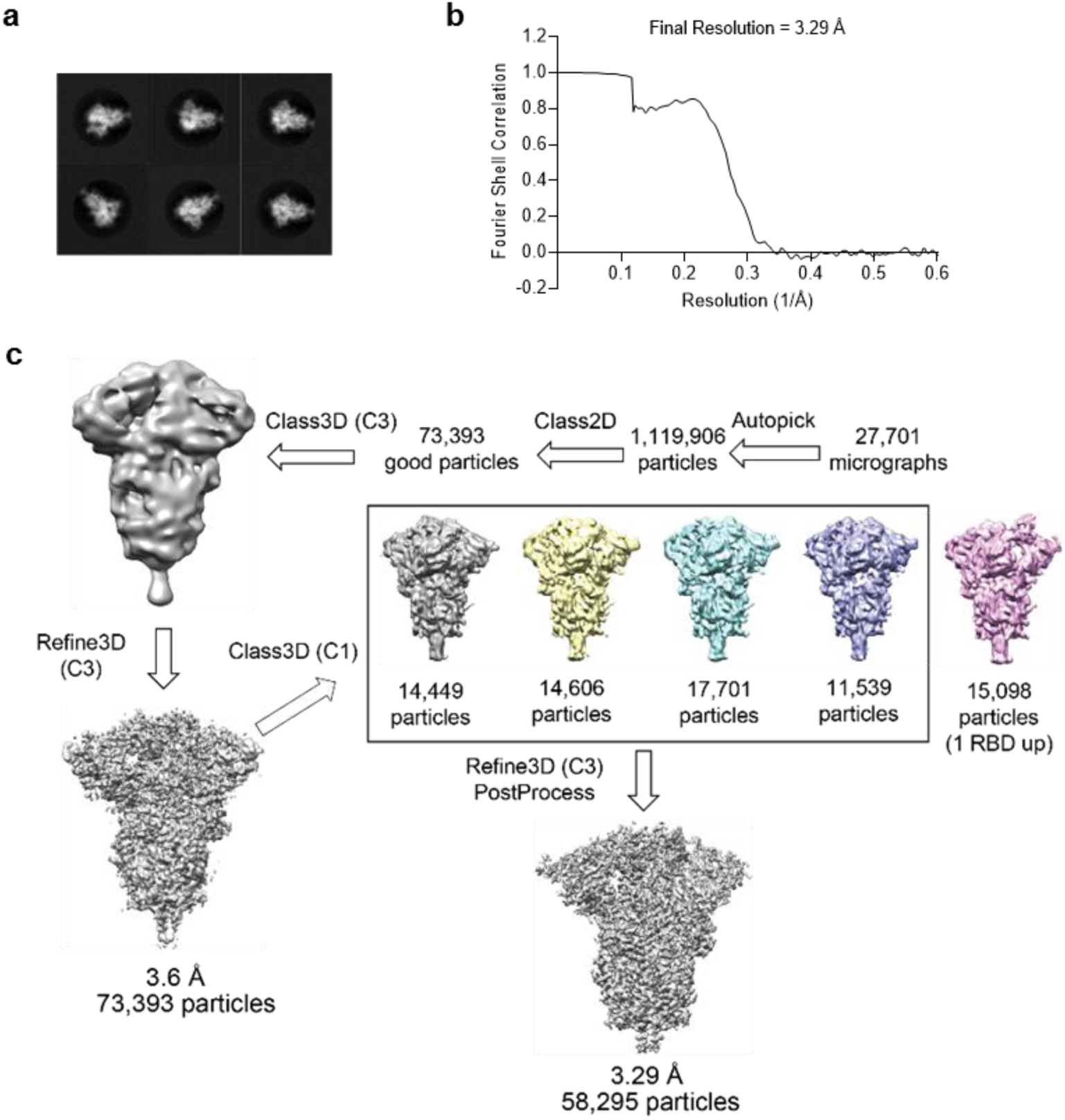
Cryo-EM evidence for alternative conformers of P2 S. **a,** Representative 2D class averages of TwinStrep-tagged P2 S particles extracted from cryo-EM micrographs. Box edge: 39.2 nm. **b,** Fourier shell correlation curve from RELION gold-standard refinement of the P2 S trimer. **c,** Flowchart for cryo-EM data processing of the complex, showing 3D class averages.

**Extended Data Figure 3.**
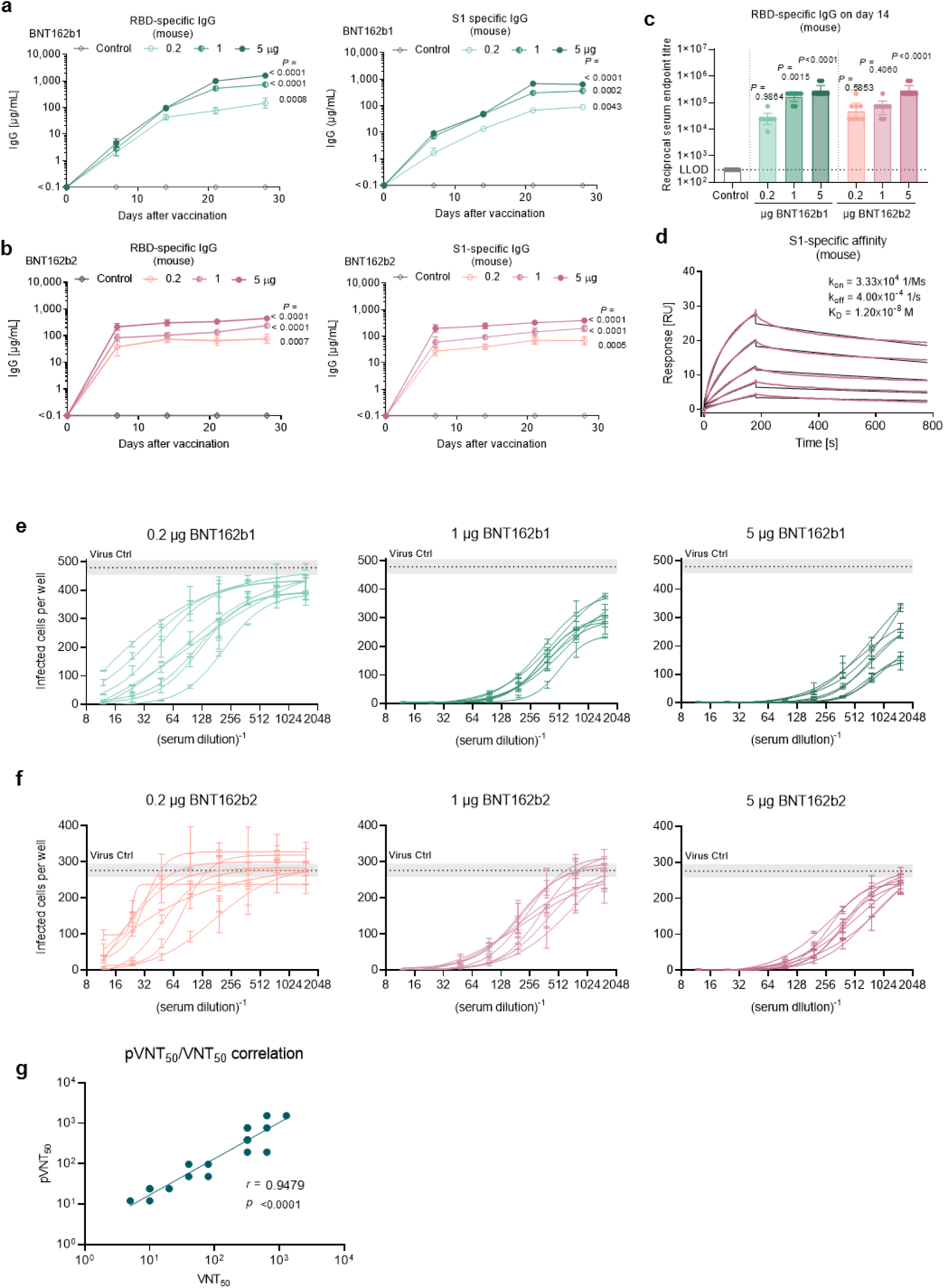
BNT162b-elicited antibody responses in mice. BALB/c mice (*n*=8) were immunised intramuscularly (IM) with a single dose of each BNT162b vaccine candidate or buffer (control, *n*=8). Geometric mean of each group (a-c) ± 95% CI (c), Day 28 p-values compared to control (multiple comparison of mixed-effect analysis [**a, b]** and one-way ANOVA [**c**], all using Dunnett’s multiple comparisons test) are provided. **a, b,** RBD- and S1-specific IgG responses in sera obtained 7, 14, 21 and 28 days after immunisation with BNT162b1 (**a**) or BNT162b2 (**b**), determined by ELISA. For day 0 values, a pre-screening of randomly selected mice was performed (*n*=4). **c,** Reciprocal serum endpoint titres of RBD-specific IgG 14 days after immunisation. The horizontal dotted line indicates the lower limit of detection (LLOD). **d**, Representative surface plasmon resonance sensorgram of the binding kinetics of His-tagged S1 to immobilised mouse IgG from serum drawn 28 days after immunisation with 5 µg BNT162b2. Binding data (in colour) and 1:1 binding model fit to the data (black) are depicted. **e, f,** Number of infected cells per well in a pseudovirus-based VSV-SARS-CoV-2 50% neutralisation assay conducted with serial dilutions of mouse serum samples drawn 28 days after immunisation with BNT162b1 (**e**) or BNT162b2 (**f**). Lines represent individual sera. Horizontal dotted lines indicate geometric mean ± 95% CI (as grey area) of infected cells in the absence of mouse serum (virus positive control). **g**, Pearson correlation of pseudovirus-based VSV-SARS-CoV-2 50% neutralisation titres with live SARS-CoV-2 virus neutralisation titres for *n* = 10 random selected serum samples from mice immunised with BNT162b1 and BNT162b2 each.

**Extended Data Figure 4.**
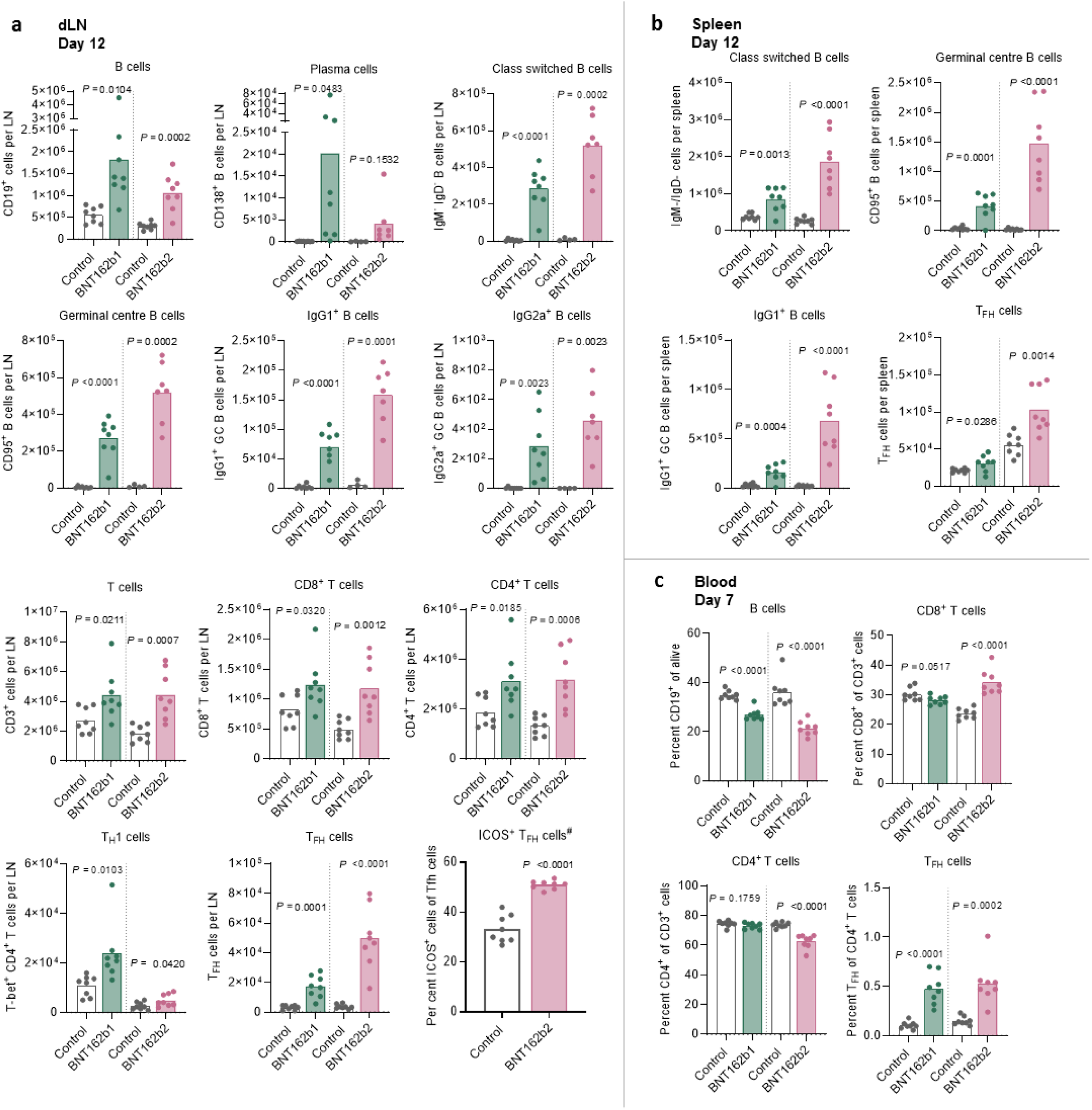
B-cell and T-cell phenotyping in lymphoid compartments of BNT162b vaccine immunised mice. BALB/C mice (*n*=8 per group) were immunised with 5 µg of each BNT162b vaccines or buffer (control). Cell subset composition was determined by flow cytometry. P-values were determined by a two-tailed unpaired t-test. **a**, B-cell and T-cell numbers in draining lymph nodes (popliteal, iliac and inguinal lymph nodes; dLN) (for B-cell subtyping: control, *n*=4, BNT162b2, *n*=7). For percent ICOS^+^ cells of TFH, only BNT162b2 data are available. **b**, B-cell and TFH-cell numbers in the spleen. **c**, B-cell and T-cell numbers in the blood.

**Extended Data Figure 5.**
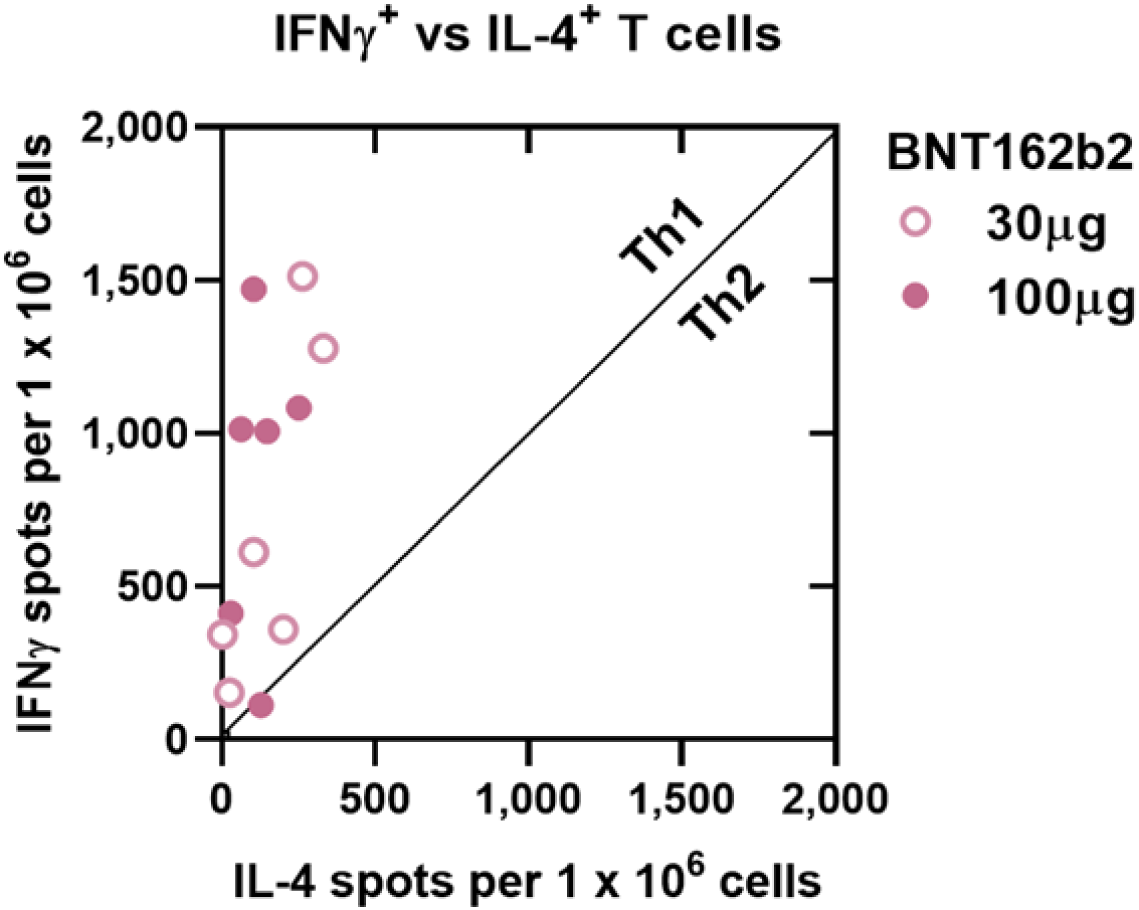
Scatterplot of IL-4 vs. IFNγ ELISpot of PBMCs from rhesus macaques immunised with BNT162b2. Rhesus macaques (*n*=6 per group) were immunised on Days 0 and 21 with 30 µg or 100 µg BNT162b2 as in Figs. 4 and 5. PBMCs for ELISpot were obtained on day 42 and were stimulated with a full-length overlapping S peptide pool. Correlation of IL-4 and IFNγ spots per 1 x 10^6^ cells.

**Extended Data Figure 6.**
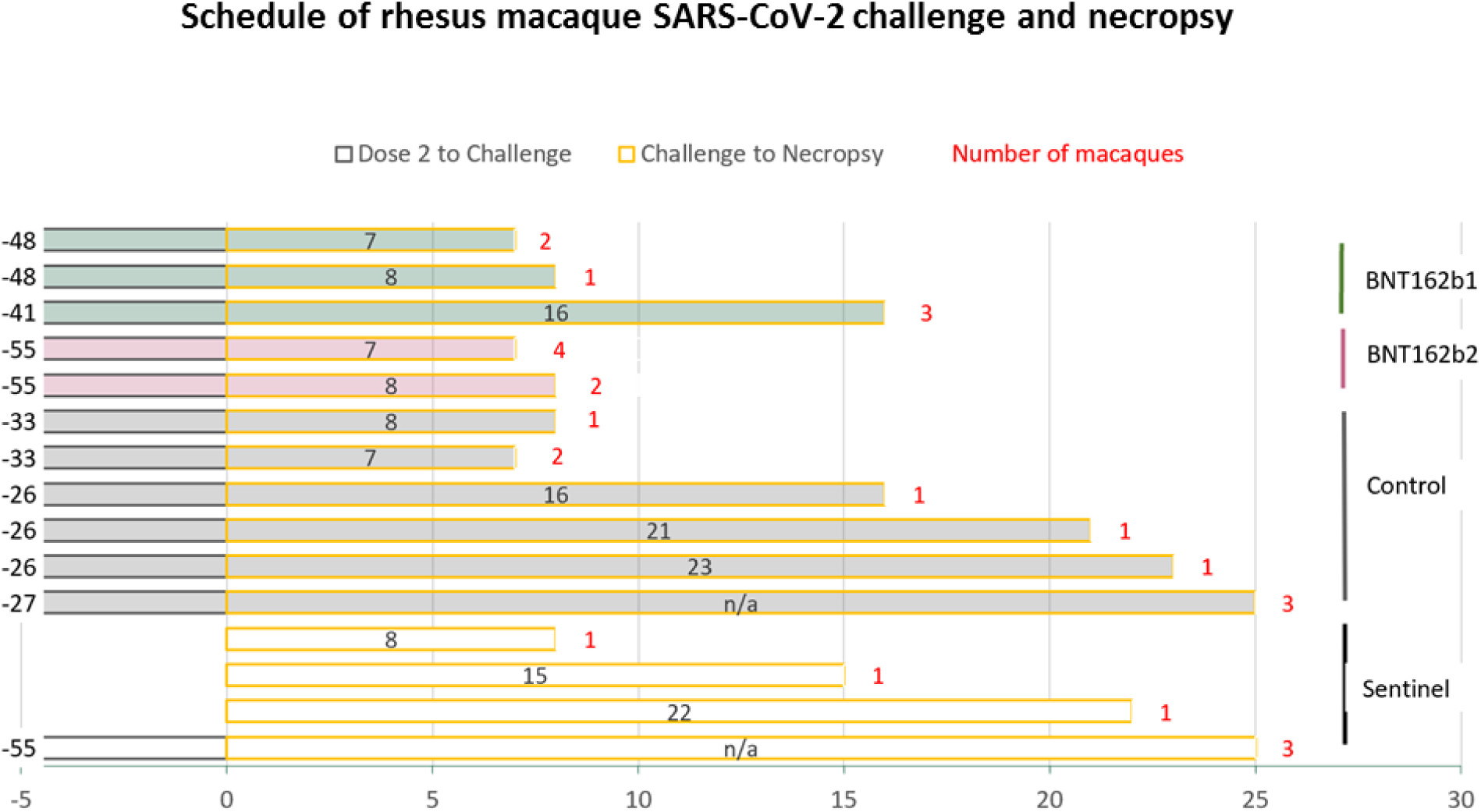
Schedule of rhesus macaque challenge and necropsy. Timing in days from Dose 2 of vaccine or saline (numbers to the left of the bars) and of necropsy (numbers inside bars) are presented relative to the day of SARS-CoV-2 or mock challenge (Day 0). Numbers of macaques represented by the bars are indicated by red numbers to the right of the bars. Control: macaques challenged but not immunised with BNT162b. Sentinel: macaques mock challenged (cell culture medium only). n/a: macaques not necropsied. Additional details, including timing of sample collections and radiographic examinations, are in Extended Data Table 2.

**Extended Data Figure 7.**
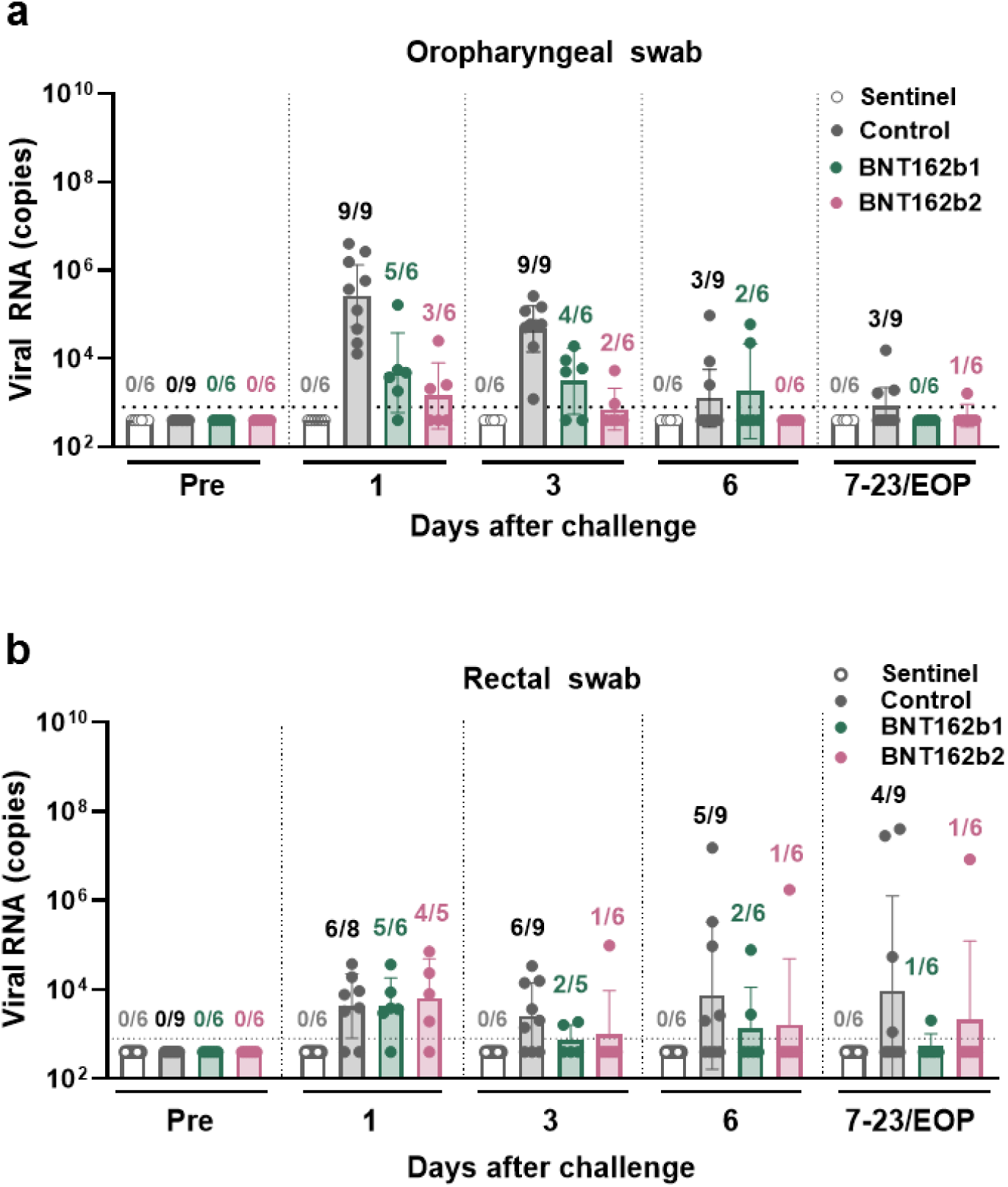
Viral RNA detection in oropharyngeal (OP) and rectal swabs from rhesus macaques after BNT162b immunisation and challenge with infectious SARS-CoV-2. Rhesus macaques immunised with 100 µg of BNT162b1 or BNT162b2 (*n*=6 each) and macaques immunised with saline or not immunised (Control, *n*=9), as described in Fig. 4, Extended Data Fig. 6, and Extended Data Table 2, were challenged with 1.05 × 10^6^ total plaque forming units (PFU) of SARS-CoV-2 split equally between the intranasal (IN) and intratracheal (IT) routes. Additional macaques (sentinel, *n*=6) were mock-challenged with cell culture medium. Viral RNA levels were detected by RT-qPCR. **a**, Viral RNA in OP swabs. **b**, Viral RNA in rectal swabs. Ratios above data points indicate the number of viral RNA positive animals among all animals providing evaluable samples in a group. Heights of bars indicate geometric mean of viral RNA copies; whiskers indicate geometric standard deviations. Every symbol represents one animal. Dotted lines indicate the lower limits of detection (LLODs). Values below the LLOD were set to ½ the LLOD. The statistical significance by Friedman’s non-parametric test of differences in viral RNA detection between 6 BNT162b1-immunised and 6 contemporaneously control-immunised animals (challenge cohorts 1 and 2) after challenge was p < 0.001 for OP swabs and p = 0.118 for rectal swabs; between 6 BNT162b2-immunised animals and 3 contemporaneously control-immunised animals (challenge cohort 3) after challenge, the statistical significance was p < 0.001 for OP swabs and p = 0.221 for rectal swabs.

**Extended Data Figure 8.**
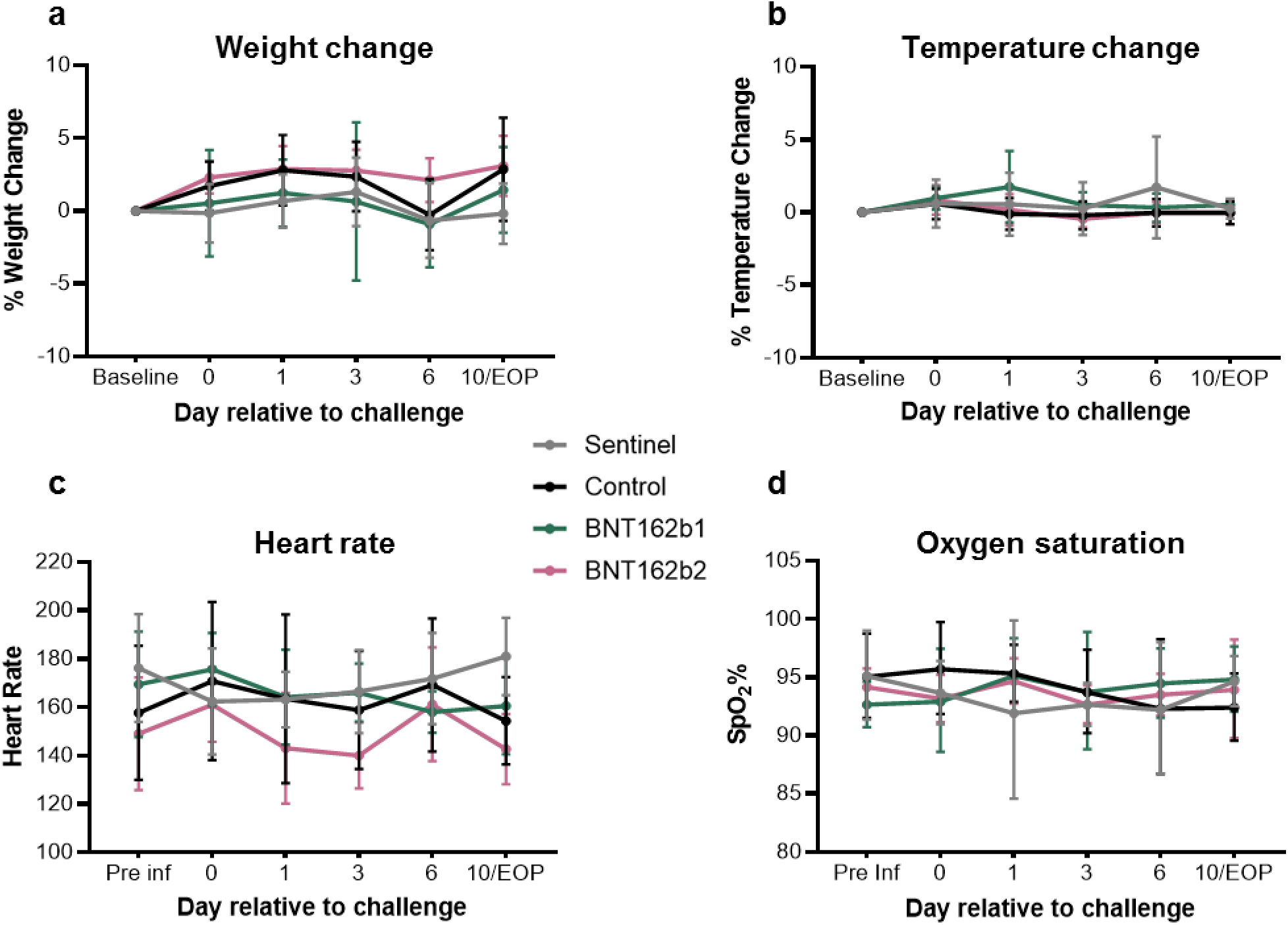
Clinical signs in BNT162b vaccine-immunised rhesus macaques after challenge with infectious SARS-CoV-2. Rhesus macaques were immunised with BNT162b vaccine candidates (*n*=6 per group) or saline (control; *n*=9) and challenged with SARS-CoV-2. A sentinel group was challenged with cell culture medium (*n*=6) as described in Figs. 4 and 5 and Extended Data Table 2. Vital signs were recorded. **a,** Body weight change. **b**, Temperature change. **c**, Heart rate. **d**, Oxygen saturation.

**Extended Data Figure 9.**
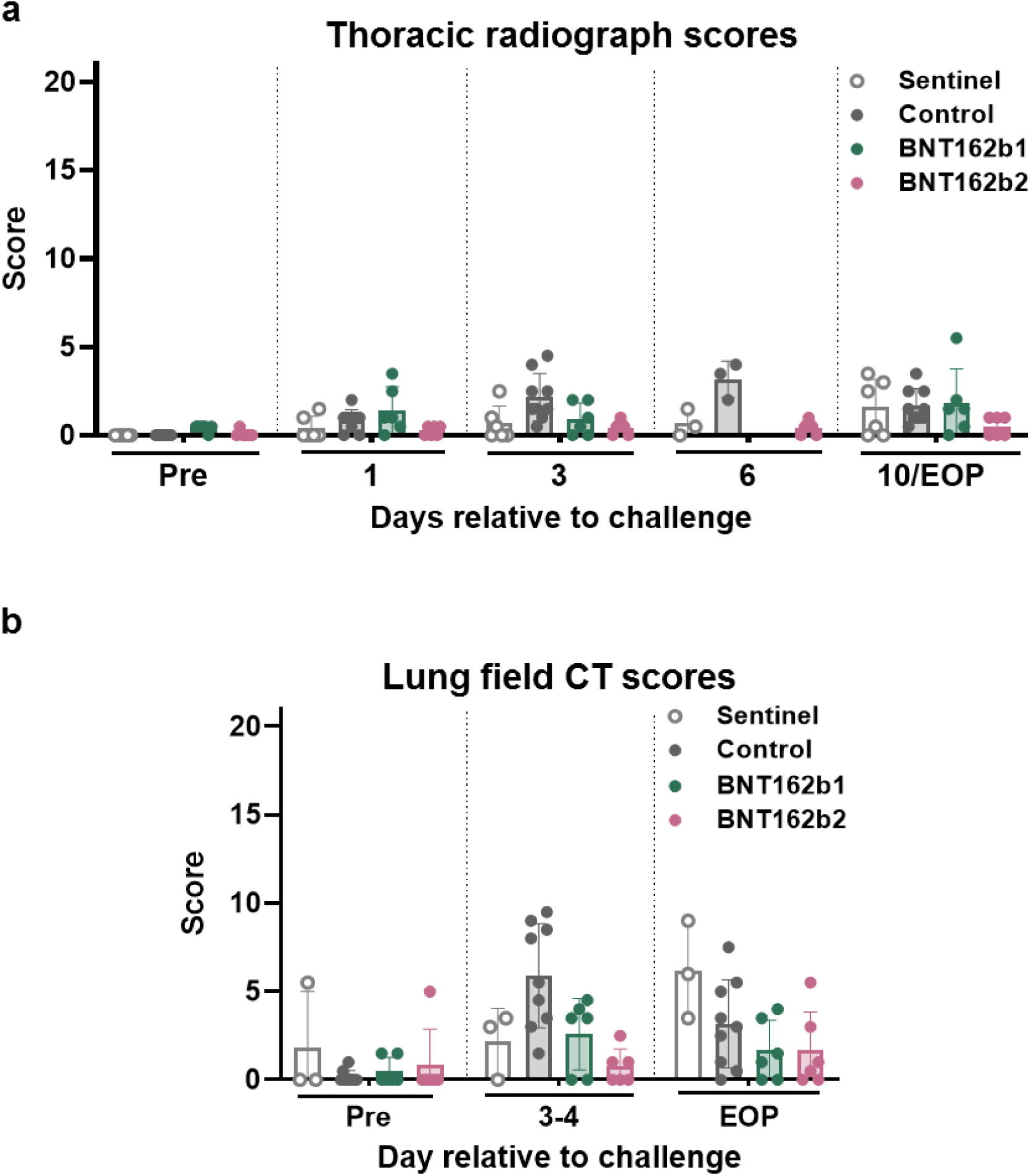
Radiographic signs in rhesus macaques after immunisation with BNT162b1 or BNT162b2 and challenge with SARS-CoV-2. Rhesus macaques were immunised with BNT162b1, BNT162b2, or saline (control) and challenged with SARS-CoV-2. A sentinel group was challenged with cell culture medium and imaged as described in Figs. 4 and 5 and Extended Data Table 2. Three-view thoracic radiographs (ventrodorsal, right and left lateral) and lung field CT images were obtained prior to challenge (pre), and post-challenge at the indicated time points. The animals were anesthetised and intubated to perform end inspiratory breath-hold. Images were interpreted by two board-certified veterinary radiologists blinded to treatment groups. Scores were assigned to 7 lung regions on a severity scale of 0-3 per region, with a maximum severity score of 21. Pulmonary lesions evident prior to challenge or those which could not be unequivocally attributed to the viral challenge (such as atelectasis secondary to recumbency and anesthesia) received a score of “0”. **a,** Thoracic radiograph scores. **b,** Lung field CT scores.

**Extended Data Figure 10.**
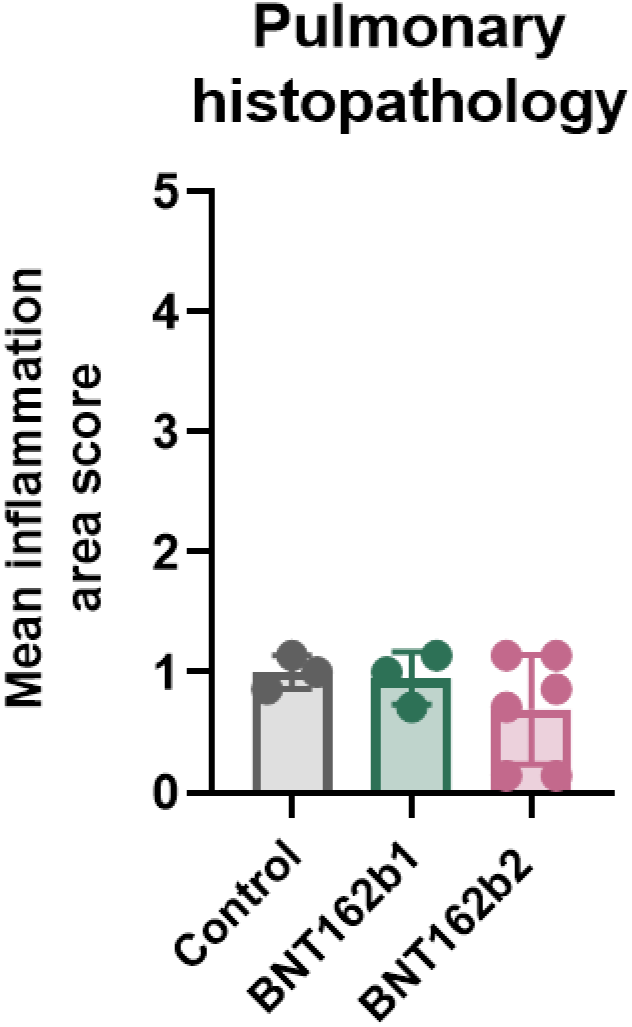
Pulmonary histopathology in rhesus macaques after immunisation with BNT162b1 or BNT162b2 and challenge with infectious SARS-CoV-2. Rhesus macaques were immunised with BNT162b1, BNT162b2, or saline (control) and challenged with SARS-CoV-2. A sentinel group was challenged with cell culture medium. The macaques were necropsied as described in Figs. 4 and 5 and Extended Data Table 2. Two veterinary pathologists blindly performed microscopic evaluation of formalin fixed, hematoxylin and eosin stained lung tissue sections from each of 7 lobes from each macaque that had been necropsied on Day 7 or 8. Inflammation scores were assigned by consensus between the pathologists on a scale of 1-5 based on the area of involvement. Each dot represents an individual animal and is the mean inflammation area score from the 7 lung lobes.

**Extended Data Table 1.**
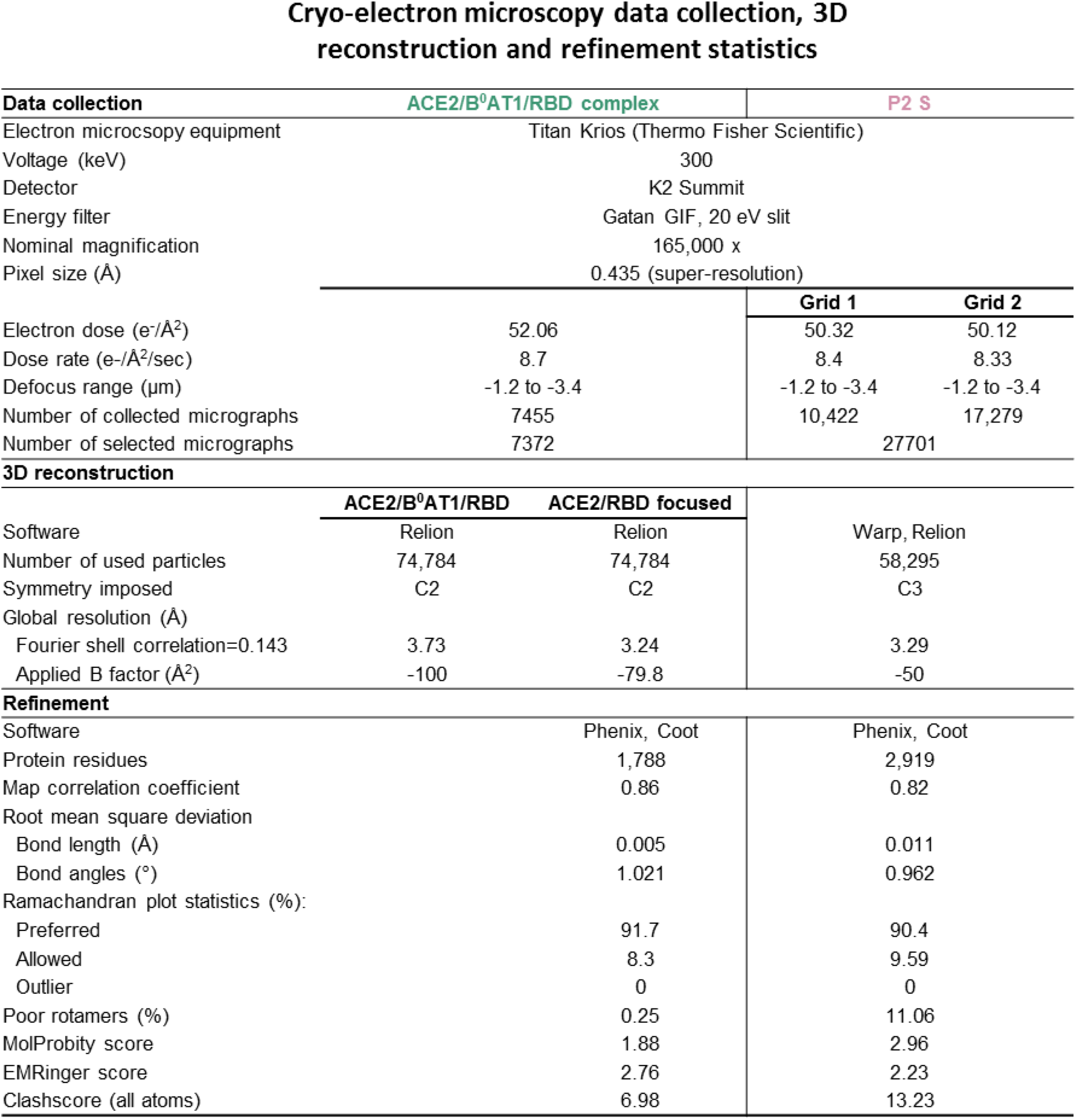
Cryo-EM data collection, 3D reconstruction and refinement statistics.

**Extended Data Table 2.**
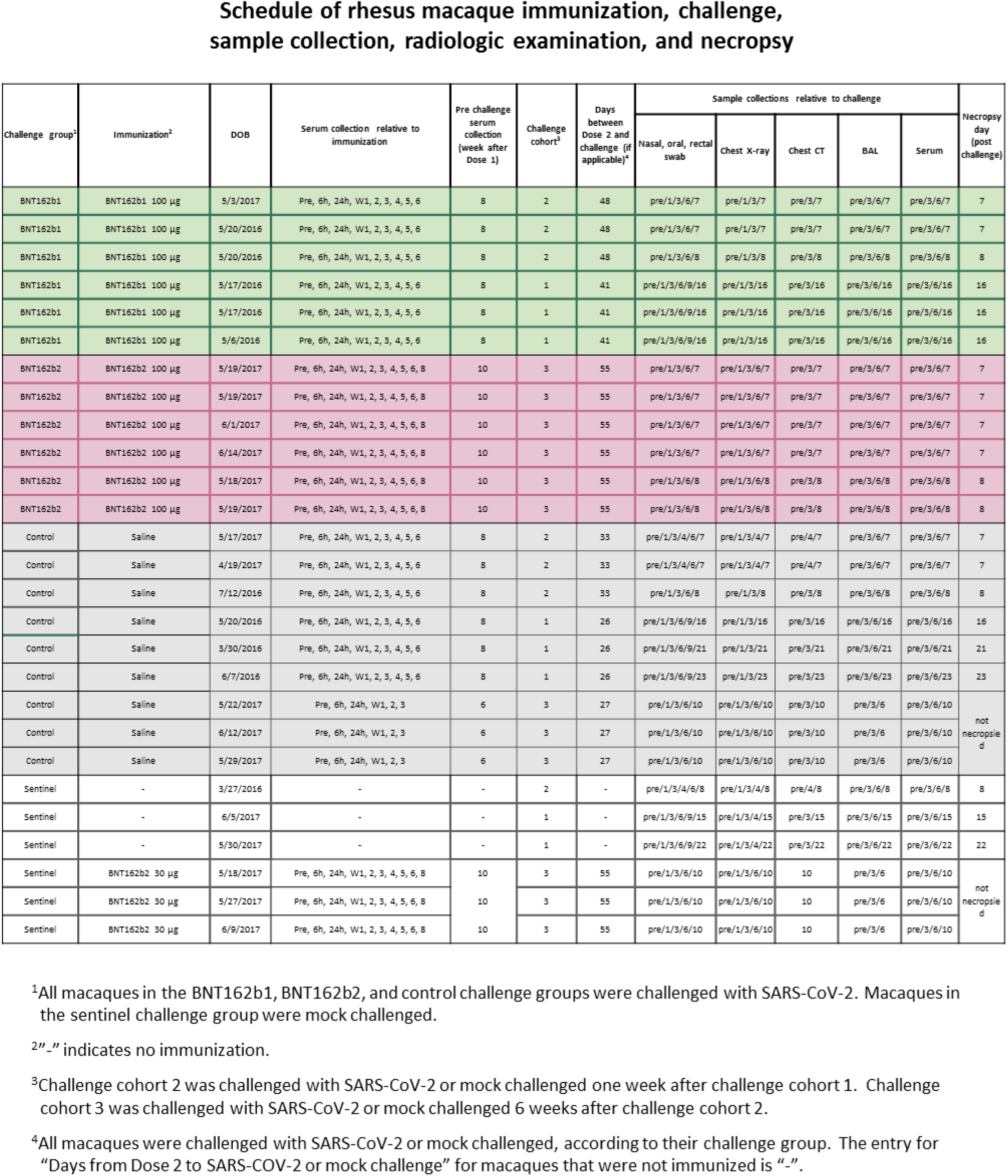
Schedule of rhesus macaque immunisation, challenge, sample collection, radiologic examination, and necropsy.

**Supplementary Figure 1.**
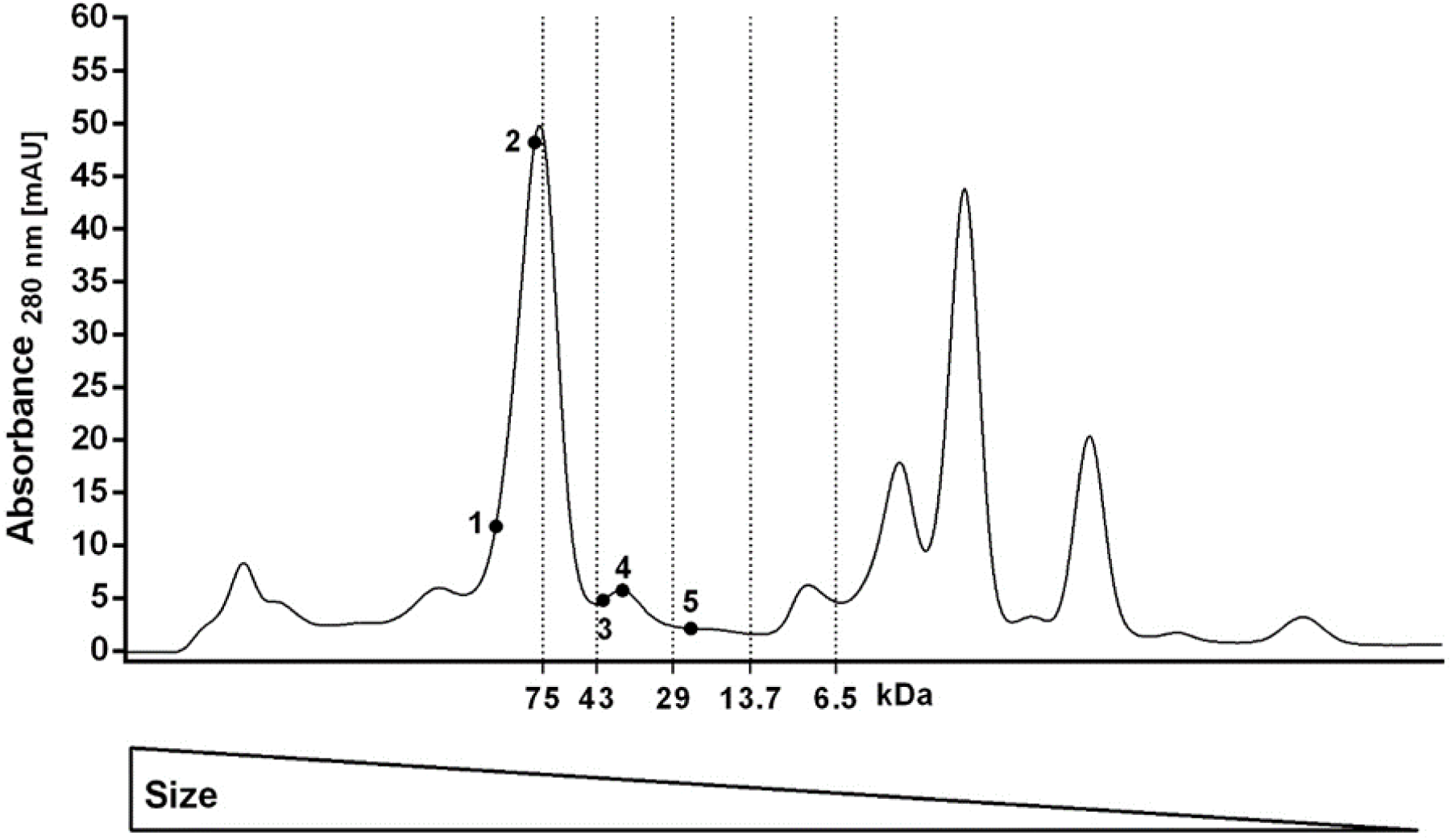
Size exclusion chromatography of medium from BNT162b1 RNA-transfected cells. Concentrated medium of HEK293T cells transfected with BNT162b1 RNA formulated with a transfection reagent (BNT162b1 RNA) was applied to a size exclusion chromatography column, calibrated using protein size standards (75, 43, 29, 13.7 and 6.5 kDa). Numbered dots indicate fractions that were further analysed by western blot.

**Supplementary Figure 2.**
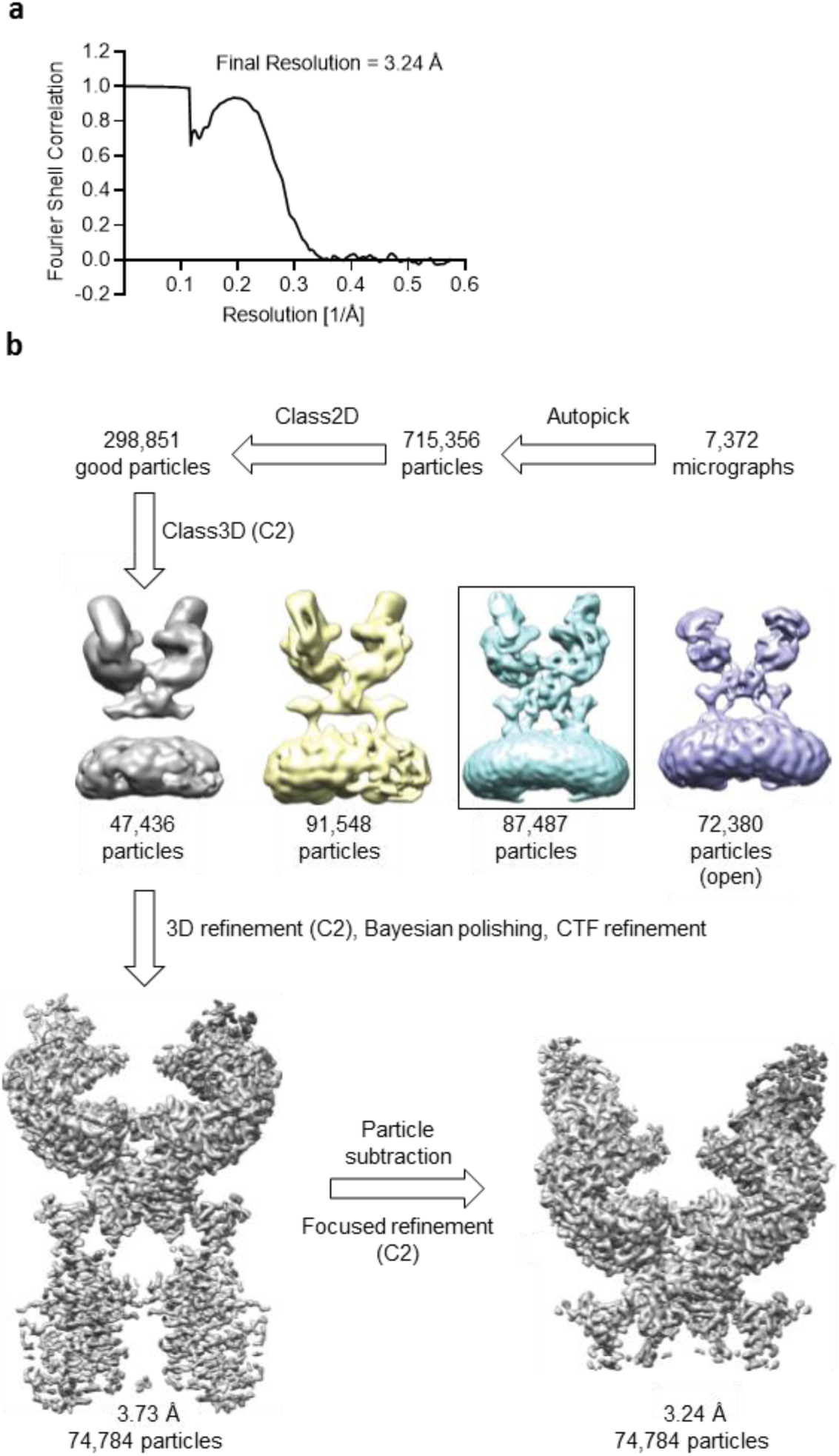
Supporting data for cryo-EM of the trimerised RBD in complex with receptor. **a**, Fourier shell correlation curve from RELION focused gold-standard refinement of the ACE2/B^0^AT1/RBD-trimer ternary complex. **b**, Flowchart for cryo-EM data processing of the complex. CTF, contrast transfer function. C2 symmetry applied during classification and refinement.

**Supplementary Figure 3.**
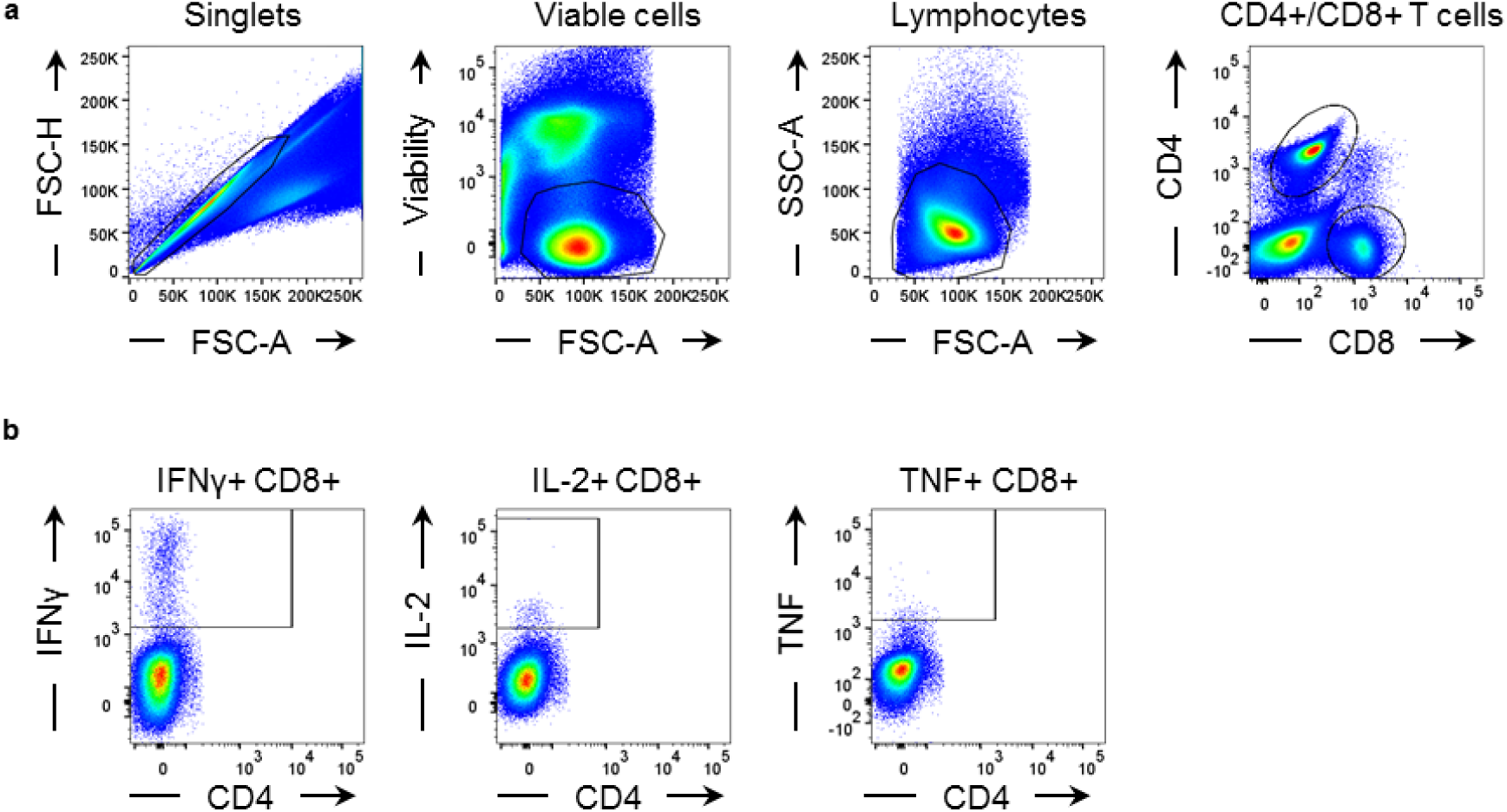
Gating strategy for flow cytometry analysis of mouse data shown in Figure 3 c. Flow cytometry gating strategy for the identification of IFNγ, IL-2, and TNF secreting CD8^+^ T cells in the mouse spleen. **a,** CD8^+^ T cells were gated within single, viable lymphocytes. **b,** Gating of IFNγ, IL-2 and TNF in CD8^+^ T cells.

**Supplementary Figure 4.**
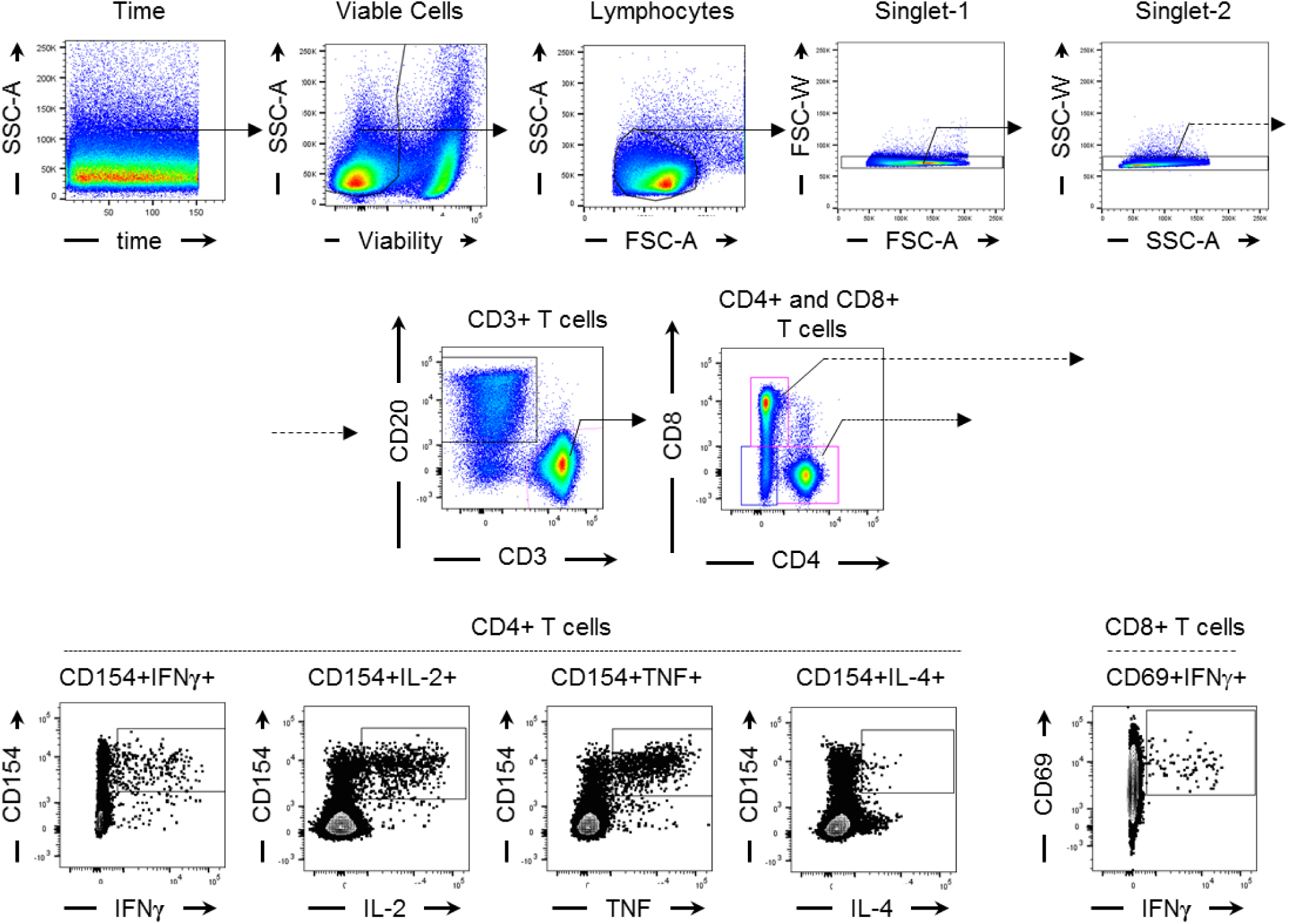
Gating strategy for rhesus macaque flow cytometry analysis of data shown in Figure 4 e-g. Flow cytometry gating strategy for identification of spike-specific SARS-CoV-2 modRNA vaccine BNT162b2-induced T cells. Starting with events acquired with a constant flow stream and fluorescence intensity, viable cells, lymphocytes and single events were identified and gated (upper row, left to right). Within singlet lymphocytes, CD20^-^ CD3^+^ T cells were identified and gated into CD4^+^ T cells and CD8+ T cells (middle row). Antigen-specific CD4+ T cells were identified by gating on CD154 and cytokine-positive cells, and CD8+ T cells were identified by gating on CD69 and cytokine-positive cells. The antigen-specific cells were used for further analysis (bottom row).

**Supplementary Figure 5.**
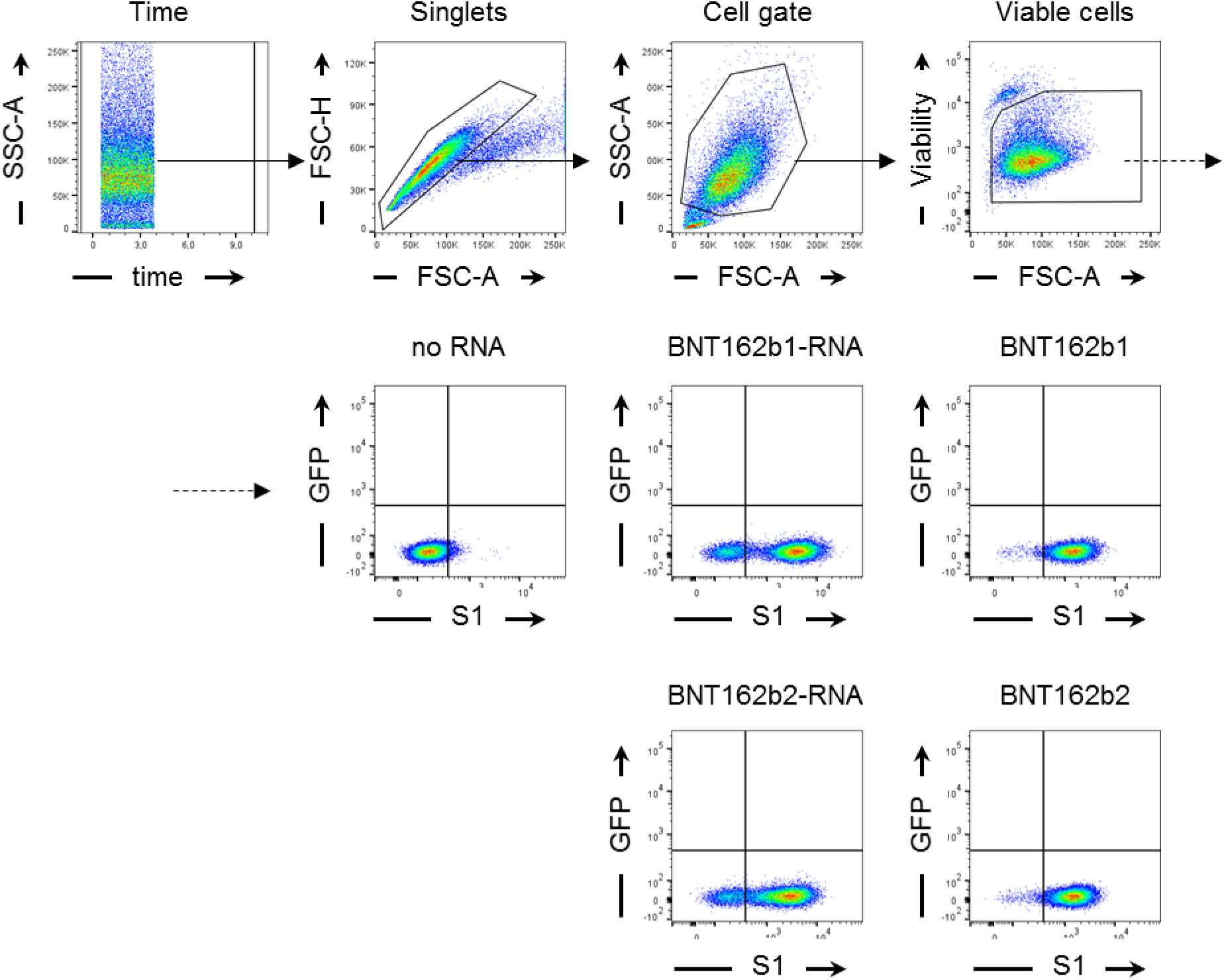
Gating strategy for flow cytometry analysis of data shown in Extended Data Figure 1a. Flow cytometry gating strategy for the identification of HEK293T cells transfected with BNT162b1 or BNT162b2, or BNT162b1-RNA or BNT162b2-RNA using a transfection reagent or no RNA (control). S1^+^ HEK293T cells were gated within single, viable HEK293T cells.

**Supplementary Figure 6.**
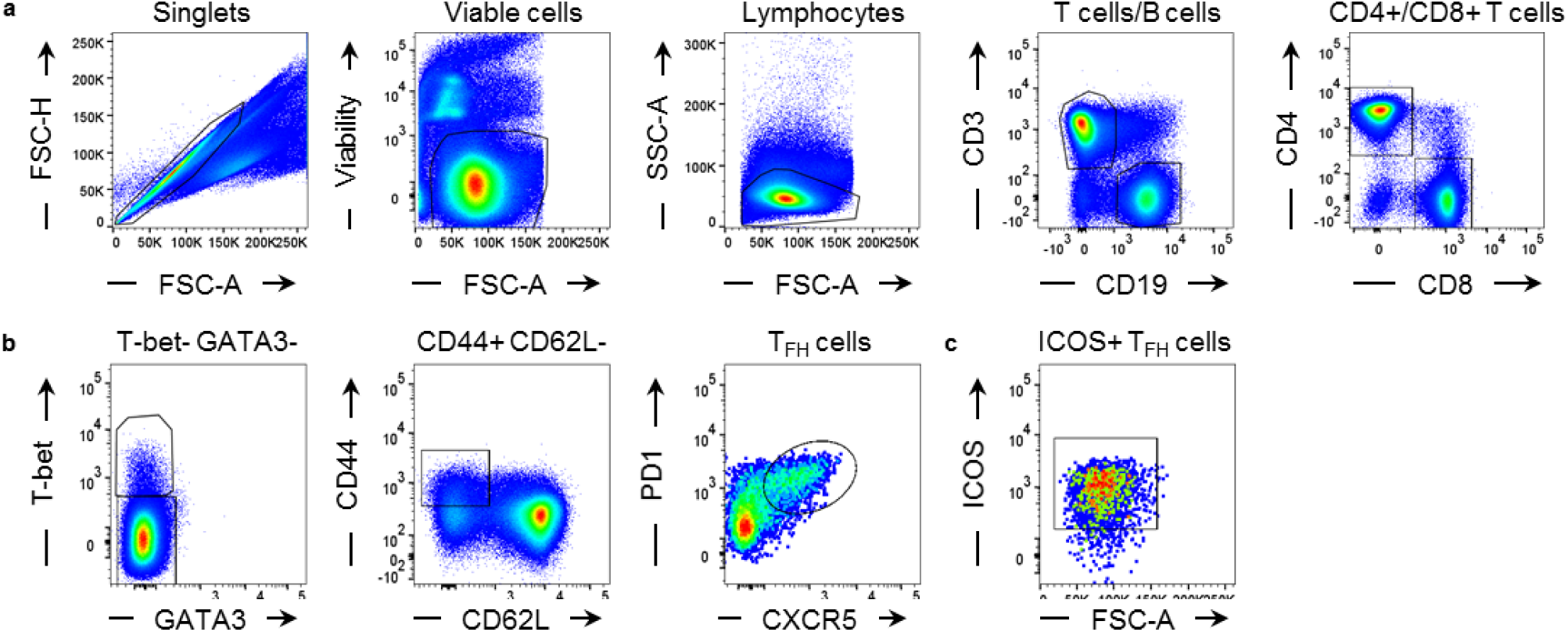
Gating strategy for flow cytometry analysis of T-cell phenotypes in murine lymph nodes and spleen shown in Extended Data Figure 4a and b. Flow cytometry gating strategy for identification of TFH cells, activated T cells and B cells in lymph nodes and the spleen. **a**, CD3^+^CD19^-^ T cells were gated within single, viable lymphocytes. CD4^+^ and CD8^+^ T cells were gated from CD3^+^ cells. **b**, TFH cells were gated from CD4^+^ T cells and defined as CD4^+^ T-bet^-^ GATA3^-^ CD44^+^ CD62L^-^ PD-1^+^ CXCR5^+^ cells.

**Supplementary Figure 7.**
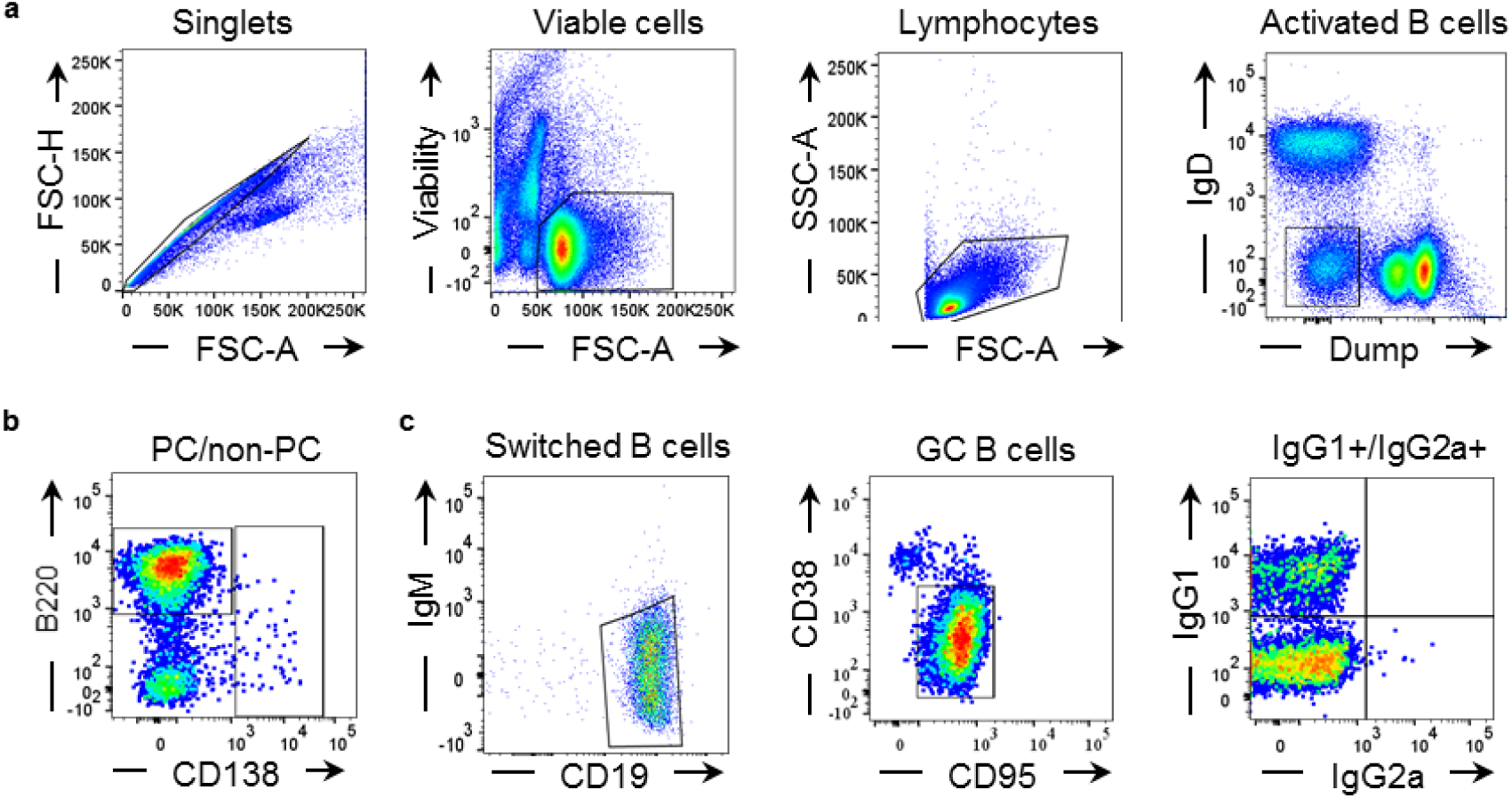
Gating strategy for flow cytometry analysis of B-cell subtypes in murine lymph nodes and spleen shown in Extended Data Figure 4a and b. Flow cytometry gating strategy for the identification of B cells in lymph nodes and the spleen. **a,** Activated B cells were gated within single, viable lymphocytes and defined as IgD-Dump (CD4, CD8, F4/80, GR-1)^-^ cells. **b,** Plasma cells (PC) were gated from activated B cells and defined as CD138^+^ B220^low/-^ cells. **c,** Switched B cells were gated from non-PC and defined as CD19^+^ CD138^-^ IgM^-^. Germinal centre (GC) and IgG1^+^ and IgG2a^+^ B cells were gated from switched B cells and defined as CD19^+^ IgM^-^ CD38^-^ CD95^+^ and CD19^+^ IgM-IgG1^+^/IgG2a^+^, respectively.

**Supplementary Figure 8.**
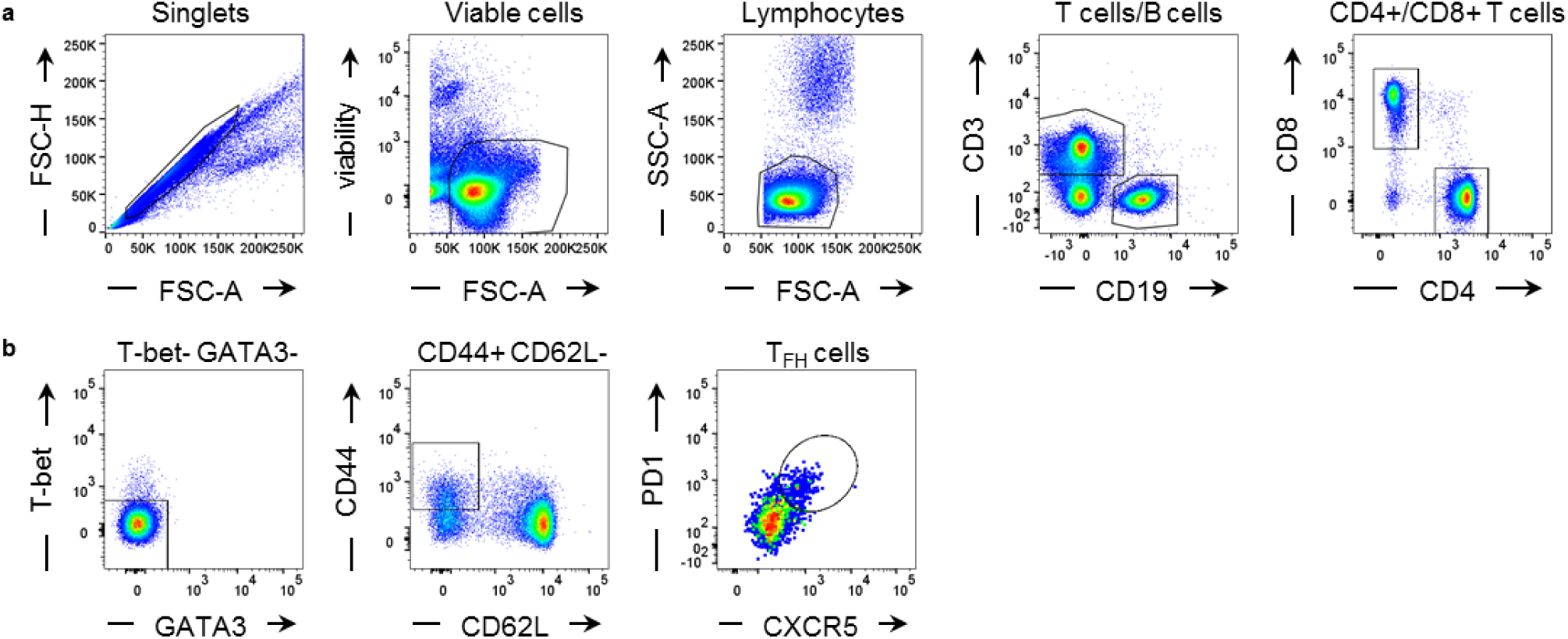
Gating strategy for flow cytometry analysis of T-cell phenotypes in mouse peripheral blood shown in Extended Data Figure 4c. Flow cytometry gating strategy for the identification of T cells, B cells and TFH cells in peripheral blood. **a,** CD3^+^ CD19^-^ T cells were gated within single, viable lymphocytes. CD4^+^ and CD8^+^ T cells were gated from CD3^+^ CD19^-^ cells. **b,** TFH cells were gated from CD4^+^ T cells and defined as CD4^+^ T-bet^-^ GATA3^-^ CD44^+^ CD62L^-^ PD-1^+^ CXCR5^+^ cells.

